# Pharmacophore-driven antibody discovery on the yeast surface

**DOI:** 10.1101/2025.10.16.682880

**Authors:** Manjie Huang, Sean J. Williams, Vikas D. Trivedi, Nikhil U. Nair, James A. Van Deventer

## Abstract

Protein-small molecule hybrids are structures capable of combining the inhibitory properties of small molecules and the specificities of binding proteins. However, discovery of such synergistic conjugates is a substantial engineering challenge. Here, we describe pharmacophore-driven antibody discovery as a high throughput approach to hybrid discovery. In this approach, we use a yeast display antibody library containing reactive noncanonical amino acids (ncAAs) and further diversify it by conjugating the library to four sulfonamide pharmacophores. Yeast display binding screens with each of the resulting billion-member hybrid collections against bovine carbonic anhydrase (bCA) yielded diverse collections of hybrids. Individual hybrids exhibited double digit nanomolar binding affinities, and frequently exhibited inhibitory properties in solution, despite the fact that the screens were based solely on binding phenotypes. Deep sequencing of sorted populations revealed that enrichments were strongly pharmacophore-dependent. In particular, screens with a potent pharmacophore led to collections of hybrids varying substantially in pharmacophore attachment point and antibody sequence features, while screens with moderate or weak pharmacophores led to collections with much narrower sets of enriched attachment points and antibody sequence features. Identification of the most frequently isolated CDR-H3 sequences and clustering CDR-H3 sequences by similarity within enriched populations provided further evidence for pharmacophore-dependent sorting outcomes. Experimental binding assays in which the pharmacophore warhead used during screening was replaced by another warhead indicated that isolated clones can tolerate alternative pharmacophores, but tend to prefer the warhead used during screening. Overall, these efforts demonstrate the utility of introducing pharmacophores into antibody libraries along with several lines of evidence that screening outcomes are pharmacophore-driven. These findings advance our understanding of hybrid discovery and highlight opportunities to pursue hybrids as research tools and potential therapeutic leads.

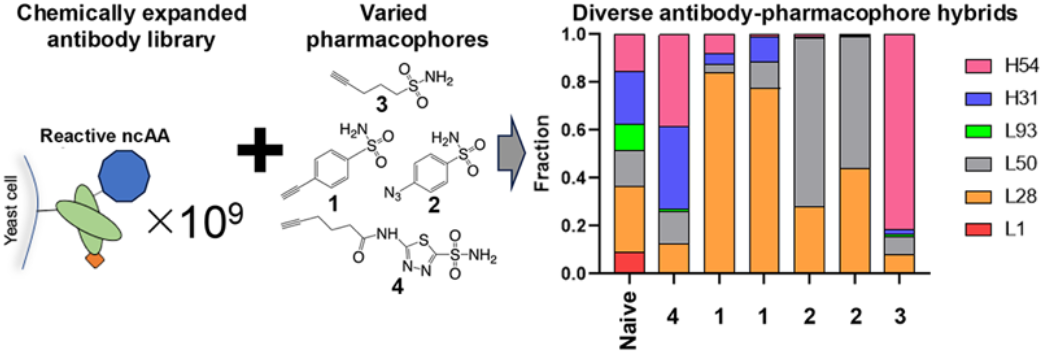

## Introduction

Despite decades of advancement in drug discovery, clinically approved inhibitors are only available for 3% of the human proteome. An additional 7% of the proteome can be addressed by inhibitors or ligands that are not clinically approved, but more than half of the proteome cannot be specifically targeted or functionally modulated.^1-3^ State-of-art tools in medicinal chemistry and drug design are now capable of dramatically improving the drug-like properties of existing hits and leads, as well as advancing new strategies for addressing difficult-to-drug targets.^3-6^ However, these impressive advances cannot overcome intrinsic limitations of small molecules: the diverse chemical functionalities of these molecules make them well-suited to bind to active sites of enzymes and other target proteins,^7^ but their limited surface areas makes it difficult for these molecules to possess single-target specificity.^8^ In contrast, the large binding surface areas of antibodies and other protein binding agents routinely exhibit high potency and specificity for a single target. Numerous discovery and engineering strategies have enabled the widespread application of binding proteins in therapeutic and diagnostic settings.^9-14^ In some cases, engineering strategies have even yielded antibodies capable of inhibiting enzymes or otherwise functionally modulating challenging targets with high specificities.^14-19^ Despite substantial advances,^20^ since most high throughput binding protein screens are limited to binding phenotypes, protein-based inhibitors remain challenging to discover and engineer.^17, 21-25^

“Hybrids” that include both protein and small molecule components offer opportunities to combine the inhibitory properties of small molecules with the potencies and specificities of proteins. Prior efforts to design and engineer hybrids have led to important proofs-of-concept, including rationally designed conjugates linking kinase inhibitors to mini-protein or fibronectin scaffolds^26, 27^ to enhance kinase inhibitor specificity, and conjugation of carbonic anhydrase inhibitors to peptides resulting in nanomolar inhibitors.^28, 29^ The Hackel Lab has established large libraries of small molecule-fibronectin hybrids and screened them to engineer potent, isoform-selective inhibitors of human carbonic anhydrases.^30^ Recently, they reported that small molecule conjugation site preference varies strongly even among isoforms from the same family of proteins.^31^ Our group investigated how different linkers and pharmacophores affect the properties of fibronectin-based carbonic anhydrase inhibitors,^32^ observing that changes in the chemical elements of hybrids influence binding and inhibitory properties in multiple ways. Despite this progress, both our fundamental understanding of how to discover and engineer hybrids and the vast combinatorial protein-small molecule hybrid spaces has only been investigated in a handful of studies.^29-34^ In particular, antibody-based hybrids remain virtually unexplored, and the effects of pharmacophore properties on the hybrid discovery process are similarly unknown. There are many outstanding questions regarding how best to “chemically diversify” binding protein structures that require critical advances in both fundamental understanding and technical platforms for hybrid discovery.

In this study, we establish a strategy for conducting pharmacophore-driven antibody discovery on the yeast surface. Using our previously described Clickable CDR-H3 synthetic antibody library encoding noncanonical amino acids (ncAAs) (**Figure 1**),^35^ we prepared several versions of the library in which pharmacophores known to inhibit carbonic anhydrases were conjugated to the billion-member library. Installing four pharmacophores of varying potencies via click chemistry enabled us to systematically screen the resulting set of hybrids against the model target bovine carbonic anhydrase (bCA). Deep sequencing revealed that distinct collections of antibody sequences emerged from screens performed with the different pharmacophores. The divergence of these populations suggests that pharmacophores strongly influence the sorting process, shifting enrichments towards sets of antibody sequences that synergize with (or at least tolerate) the pharmacophore installed in the library. Characterization of individual hybrid clones of interest on the yeast surface revealed that several hybrid clones exhibit apparent K_D_s in the double-digit nanomolar range, even when the pharmacophore is known to exhibit micromolar-range binding. Removal of pharmacophores from hybrids led to undetectable levels of bCA binding in all clones evaluated in this work. In addition, “warhead swap” experiments in which the pharmacophore used during sorting was replaced with other pharmacophores commonly resulted in reduced bCA binding, further indicating the key role of the pharmacophore in combination with the antibody paratope in hybrid structures. Finally, preparation of soluble forms of hybrids revealed that several hybrids exhibit improved inhibitory properties compared to pharmacophores alone in standard biochemical assays, even though our screens were based solely on target binding as a screening criteria. Overall, these findings establish pharmacophore-driven antibody discovery on the yeast surface as a powerful discovery platform and demonstrate the critical role of the pharmacophore in shaping screening outputs. Efficient explorations of hybrid sequence spaces provide opportunities for the discovery of unique tools for advancing basic research, diagnostics, and therapeutics.

**Figure 1.**
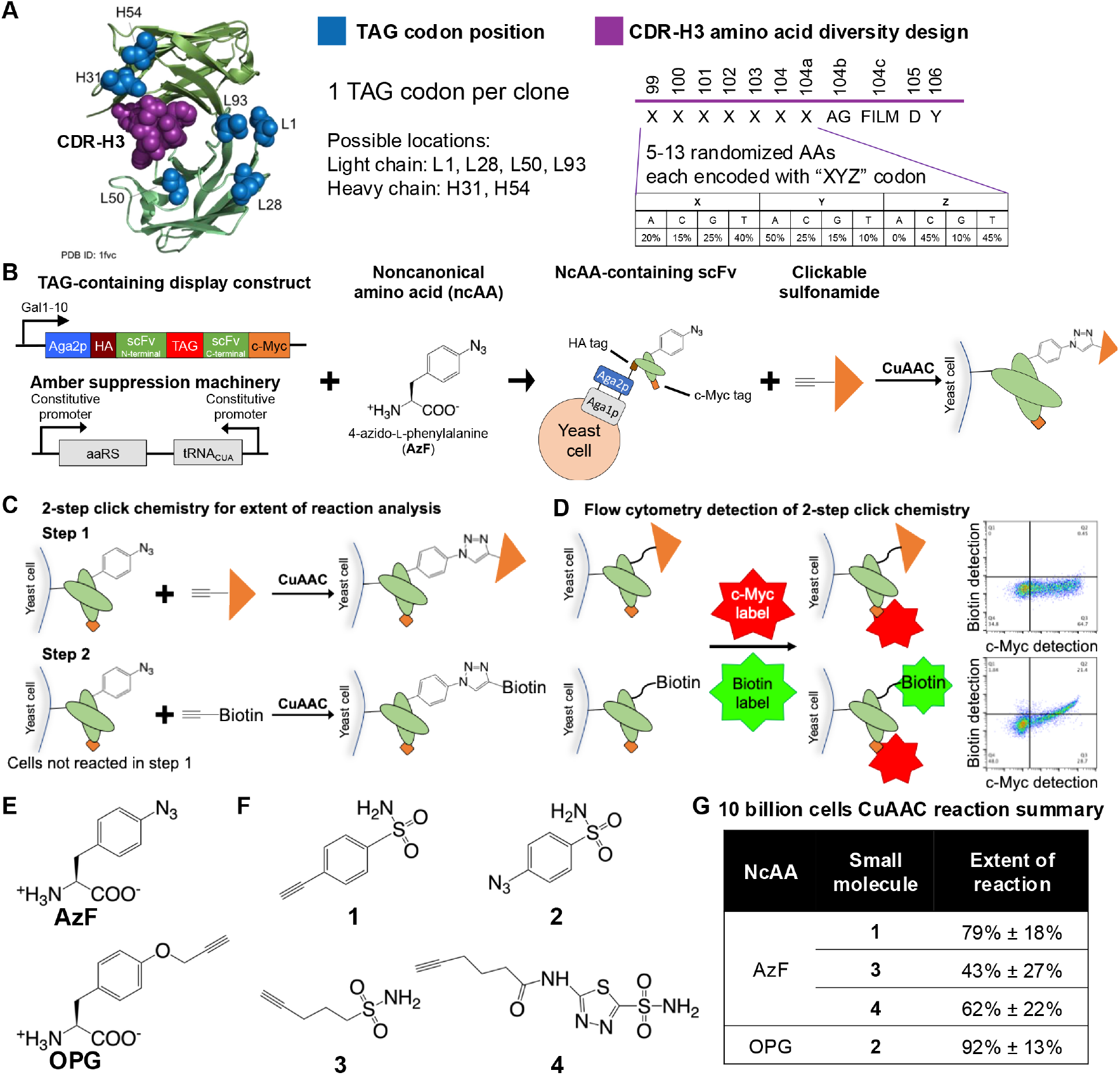
Construction of antibody-small molecule hybrids on the yeast surface. **A)** Design of the clickable CDR-H3 library. The 1 billion different clones each contain a TAG codon at one of the six positions (blue) and a diversified CDR-H3 loop (purple) that ranges from 9 to 17 amino acids in length. The diversified amino acids “X” are encoded with “XYZ” codons that mimic the amino acid frequencies found within antibodies.^35, 39^ **B)** Each yeast cell in the library is transformed with a plasmid encoding a yeast display version of an scFv variant and a separate plasmid encoding the orthogonal translation system (OTS). Clickable noncanonical amino acids (ncAAs) are incorporated at the site of the stop codon, adding a 21st amino acid to these constructs. Small molecules are then conjugated to ncAAs with copper-catalyzed azide-alkyne cycloaddition (CuAAC) chemistry. **C)** A 2-step reaction scheme facilitates validation of CuAAC click chemistry conjugation via flow cytometry. In the first step, a reaction with the cells displaying ncAA-containing scFvs is performed with a clickable small molecule. Following washing to remove excess reactants, a second CuAAC reaction is used to install a clickable biotin probe to enable detection of ncAAs that remain unreacted following Step 1 (using a fluorescent streptavidin detection probe). **D)** Fluorescent labeling of cells after 2-step click chemistry. Cells are labeled for full length display with an anti-c-Myc antibody (and fluorescent secondary antibody) and for biotin conjugation with fluorescent streptavidin. If the small molecule has reacted with all available conjugation sites, biotin will not be detected in the flow cytometry dot plot (top panel). If reactive sites remain after the reaction with the small molecule, biotin will be detected in the flow cytometry dot plot (bottom panel). **E)** Clickable ncAAs used in this work: *p*-azido-L-phenylalanine (AzF) and *O-*propargyloxy-L-phenylalanine (OPG). **F)** Clickable sulfonamides used in this work: 4-ethynyl-benzenesulfonamide (**1**); 4-azidobenzenesulfonamide (**2**); pent-4-yne-1-sulfonamide (**3**); and N-(5-sulfamoyl-1,3,4-thiadiazol-2-yl)hex-5-ynamide (**4**). **G)** Extents of reaction for initial hybrid library preparations (10 billion cells/pharmacophore). Cells were subjected to CuAAC with 1 mM small molecule pharmacophore in 25 mL for 4 hours at room temperature. Images from Figure 1A adapted from Rezhdo et al.^35^ Copyright 2025 *ACS Synthetic Biology*.

## Results

### Hybrid library preparation on the yeast surface

Previously, our lab reported the Clickable CDR-H3 library, a multi-modal, chemically diversified antibody library encoding approximately 1 billion different single-chain variable fragments (scFvs). DNA encoding each library member includes a TAG codon at one of 6 potential positions for presentation of a ncAA within the scFv as well as a diversified complementarity-determining region H3 (CDR-H3) loop (**Figure 1A, Supplementary Table S1**). In the name of the TAG codon positions, the “L” stands for light chain positions according to Kabat numbering, and “H” stands for a heavy chain position in Kabat numbering. The spatial separation of the reactive ncAA site from the diversified amino acids of CDR-H3 is distinct from previously published hybrid libraries, where encoded pharmacophore attachment points are integrated within diversified binding loops.^30, 31^ In addition to the display construct encoding an scFv with a TAG codon, each yeast cell within the library also contains a plasmid encoding the orthogonal translation system (OTS): *p*-acetyl-phenylalanyl-tRNA synthetase (TyrAcFRS) and its corresponding tRNA _CUA_^Tyr^.^36-38^ This enables hybrid construction on the yeast surface by 1) incorporating clickable ncAAs into scFv variants in the library; and 2) use of copper-catalyzed azide-alkyne cycloaddition (CuAAC) to conjugate small molecules to the scFvs (**Figure 1B**).^39^ Conjugation efficiencies are evaluated using our established 2-step approach for conjugations on the yeast surface (**Figure 1C**).^39, 40^ In the first step, cells displaying scFvs are subjected to CuAAC with small molecules and washed. In the second step, cells are then subjected to CuAAC with a biotin probe to detect the presence of ncAAs that did not react with the small molecule. After additional washing, cells are labeled to detect biotin and C-terminal tags to determine the extent of small molecule reaction (**Figure 1D**).

We prepared four different forms of the library using two reactive ncAAs and four pharmacophores. To prepare “clickable” forms of the library, we induced yeast display with the library under conditions that enabled incorporation of clickable ncAAs that are translationally active with TyrAcFRS: *p*-azido-L-phenylalanine (AzF) and *O*-propargyloxy-L-phenylalanine (OPG) (**Figure 1E**). We then introduced pharmacophores via CuAAC. Here, we used sulfonamide pharmacophores to target the active site of bCA, a well-behaved model target. We identified four commercially available, clickable sulfonamides for conjugation with the library (**Figure 1F**): *N*-(5-sulfamoyl-1,3,4-thiadiazol-2-yl)hex-5-ynamide (**4**, K_i_ ∼10 nM) contains the highly potent acetazolamide (AAZ) pharmacophore,^30^ 4-ethynyl-benzenesulfonamide (**1**, K_i_ ∼100-1000 nM) and 4-azidobenzenesulfonamide (**2**, K_i_ ∼100-1000 nM) are clickable benzenesulfonamides (BS) with moderate potency, and pent-4-yne-1-sulfonamide (**3**, K_i_ >10,000 nM) is an alkyl sulfonamide (AS) with further reduced potency.^41-43^ To preserve the diversity of the scFv library, we conducted large-scale CuAAC with 10 billion cells (10× the number of transformants in the library) with each of the four clickable small molecules. Extent of reaction analysis indicated that the majority of displayed antibodies in each 10-fold library oversampling were modified after reaction with small molecule pharmacophores **1, 2**, or **4**, and approximately half of displayed antibodies were modified after reaction with small molecule pharmacophore **3** (**Figure 1G**). Overall, we were successful in generating four hybrid forms of our billion-member, multi-modal library for high throughput screening.

### Pharmacophore-driven library sorting

After the construction of the hybrid collections, we used a series of magnetic bead-based enrichments and fluorescence-activated cell sorting (FACS) in an attempt to isolate clones capable of binding to bCA (**Figure 2, Supplementary Figure S1**).^39, 44^ We chose bCA as a target because it is both well-behaved and frequently used when establishing new drug discovery strategies, including several prior hybrid peptide studies.^28, 29, 33, 34, 45, 46^ Since each round of screening involved the installation of pharmacophores within displayed proteins on the surfaces of freshly induced cells, we evaluated the extent of reaction for each population to be sorted (**Supplementary Table S2**). As a control for pharmacophore libraries, we also attempted to enrich libraries containing only ncAAs without pharmacophore installation. We monitored sorting progress by evaluating bCA binding to each pharmacophore-modified and control population (**Supplementary Figure S1**). For all pharmacophores, 1-2 rounds of bead-based enrichments were sufficient to detect populations of bCA-binding cells during flow cytometry analysis. In contrast, the no-pharmacophore library preparations showed no evidence for enrichment under the same conditions (**Supplementary Figure S1**).

**Figure 2.**
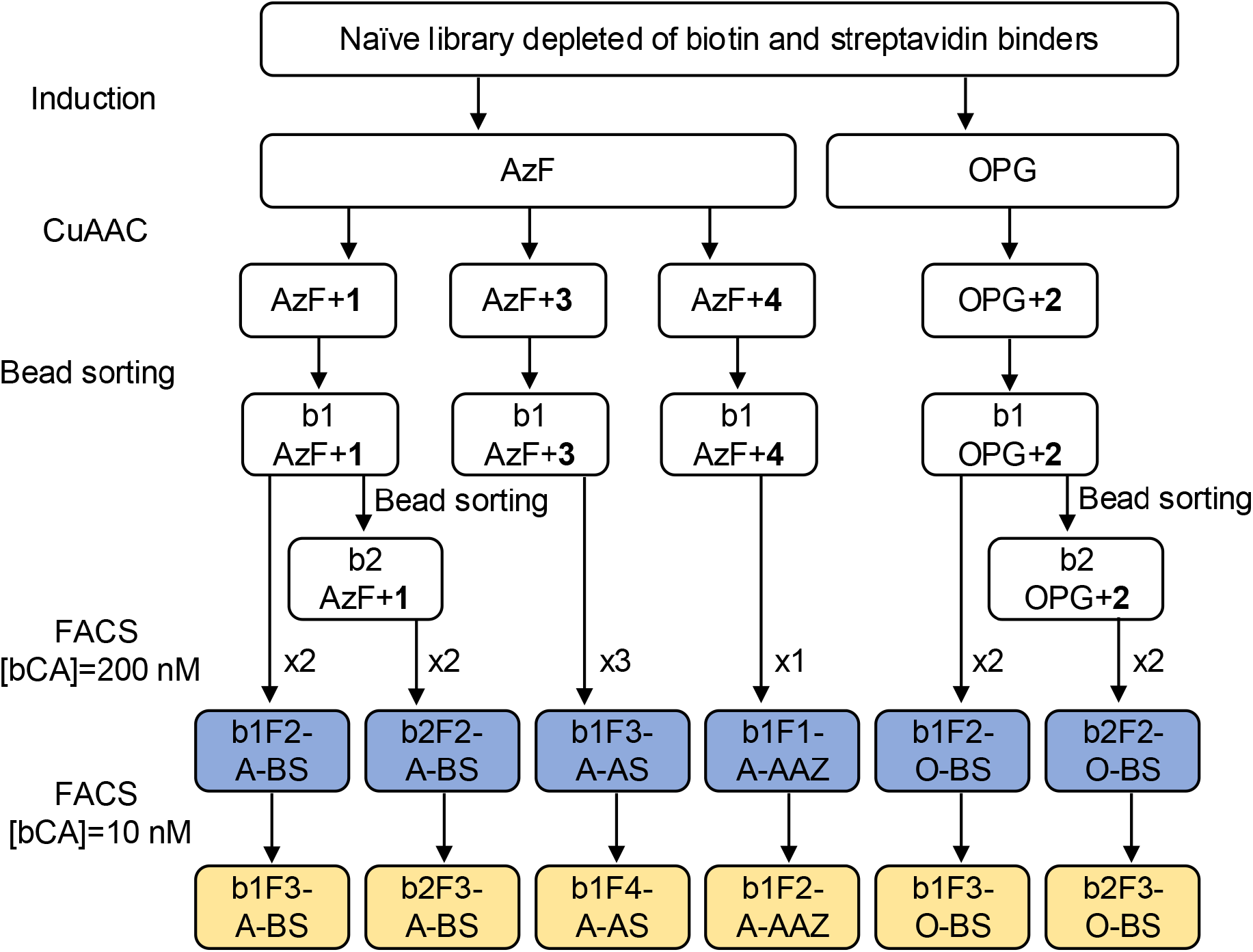
Sorting process of hybrid populations containing different pharmacophores. In the population names, bx = x rounds of bead sorting, Fx = x rounds of fluorescent activated cell sorting (FACS), A = AzF, O = OPG, BS = benzenesulfonamide (**1** or **2, 1** only reacts with AzF and **2** only reacts with OPG), AS = Alkyl sulfonamide (**3**), and AAZ= acetazolamide (**4**). Initial rounds of FACS were performed at 200 nM bovine carbonic anhydrase (bCA), and for the last round of FACS the bCA concentration was reduced to 10 nM to attempt to isolate higher affinity clones. The populations highlighted in blue were analyzed with deep sequencing; individual clones were isolated from the populations highlighted in yellow.

Following bead-based enrichments, we conducted multiple rounds of FACS to further enrich for bCA binders. Initial rounds were performed after treating hybrid-displaying cells with 200 nM bCA, while for the final round of FACS for all pharmacophores, bCA concentration was reduced to 10 nM in an attempt to enrich for higher affinity clones. The number of rounds of bead enrichment and FACS varied from pharmacophore to pharmacophore. For example, the AAZ-modified population showed the highest frequency of binding clones to bCA after 1 round of bead sorting and 1 round of FACS; this is consistent with the high potency of AAZ. In contrast, the benzenesulfonamide-modified populations required 1-2 rounds of bead enrichments followed by 2 rounds of FACS to obtain enriched populations exhibiting high bCA-binding signals. Screens of the alkyl sulfonamide-modified population required 1 round of bead enrichment and 3 rounds of FACS prior to exhibiting a population with a high frequency of antigen binding. Populations displaying ncAAs alone did not show detectable levels of bCA binding after 1 round of bead-based enrichment and 1 round of FACS enrichment and were not carried forward further. The observation that only pharmacophore-modified forms of the library led to enrichment of binding clones under the sorting conditions used here strongly suggests that installation of the small molecules influences the sorting process.

To further investigate the role of the pharmacophores in library screening, we performed deep sequencing analysis of naïve and sorted library populations (**Figure 3**; b1F2-A-BS, b2F2-A-BS, b1F3-A-AS, b1F1-A-AAZ, b1F2-O-BS, and b1F2-O-BS; see

**Figure 3.**
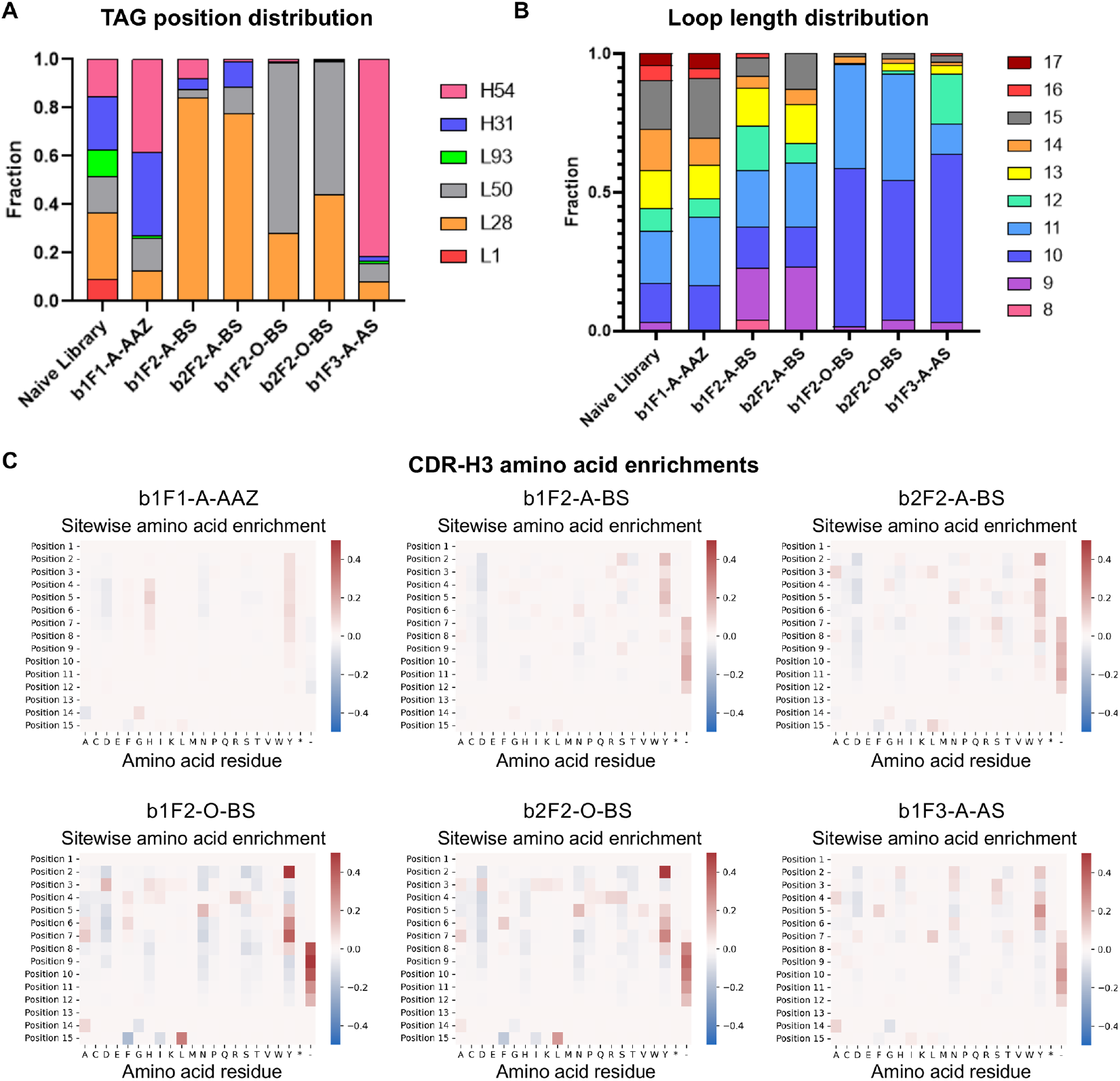
Deep sequencing analysis of the populations sorted with different pharmacophores. **A)** Distribution of TAG positions in the naïve library and in populations sorted with different pharmacophores. **B)** Distribution of loop lengths in the naïve library and in populations sorted with different pharmacophores. **C)** Site-specific amino acid enrichment heatmaps from ScaffoldSeq analysis (images generated using Python *pandas* and *seaborn* packages; see *Materials and Methods* for details). Position 1 corresponds to heavy chain position 98 (H98; Kabat numbering) of CDR-H3, and position 15 corresponds to the FILM-diversified position directly preceding H105 (The position denoted H104c in Figure 1).

Figure 2 caption for explanation of naming scheme). We investigated frequencies of TAG codon (ncAA + pharmacophore) positions, CDR-H3 loop lengths, and amino acid compositions of the CDR-H3 loops in naïve and enriched populations. Hybrid structures must effectively integrate a binding protein’s paratope with a small molecule in such a way that both can bind to the target. The frequency of ncAA + pharmacophore substitution site in enriched populations gives insights into which sites are preferred (or at least tolerated). The naïve library contains 6 distinct TAG codon positions with relatively evenly distributed frequencies, while enriched populations exhibit a range of apparent substitution preferences (**Figure 3A**). The sequenced AzF + AAZ (**4**) population (b1F1-A-AAZ) retains four of the six substitution sites at moderate to high frequencies. The AzF + benzenesulfonamide (**1**) populations (b1F2-A-BS and b2F2-A-BS) are strongly enriched for clones encoding TAG at position L28, while the OPG + benzenesulfonamide (**2**) populations (b1F2-O-BS and b2F2-O-BS) are strongly enriched for clones encoding either TAG position L28 or L50. Finally, the AzF + alkyl sulfonamide (**3**) population (b1F3-A-AS) is strongly enriched for clones encoding TAG at position H54. These differences in preferred TAG sites suggest that the distinct pharmacophores prefer different spatial relationships with the antibody paratope to better engage with the bCA active site. The substantial changes in TAG site frequencies highlights the value of encoding multiple ncAA substitution sites within a hybrid library.

In hybrids, the antibody paratope must be compatible with small molecule binding. In our simple synthetic antibody library architecture, only CDR-H3 loop length and sequence are allowed to vary and thus we evaluated how each of these elements changed within the enriched populations. For CDR-H3 loop length (**Figure 3B**), the enriched b1F1-A-AAZ population contains clones with loop lengths of 10-17 amino acids at similar frequencies. This is consistent with the high potency of AAZ, which is known to drive carbonic anhydrase binding on its own even if installed in a protein without any detectable binding activity toward carbonic anhydrases.^30^ The AzF + benzenesulfonamide (**1**) populations b1F2-A-BS and b2F2-A-BS contain clones with loop lengths ranging from 9-15 amino acids at substantial frequencies (a shift towards shorter loop lengths compared with the naïve library), but no apparent preference for an individual loop length. In contrast, the vast majority of clones in the OPG + benzenesulfonamide (**2**) populations b1F2-O-BS and b2F2-O-BS encode for CDR-H3 loop lengths of 10 or 11, and greater than 50 percent of the clones in the AzF + alkyl sulfonamide (**3**) population b1F3-A-AS encode for a CDR-H3 loop length of 10. These populations thus exhibit strong loop length preferences, in contrast to the other enriched populations. The pronounced differences in loop length distributions suggest that pharmacophores influence enrichment outcomes beyond just the preferred pharmacophore site.

We also investigated the amino acid sequence compositions of the CDR-H3 loops of enriched populations (**Figure 3C**; separate analyses performed on each CDR-H3 loop length provided in **Supplemental Figure S2**). Leveraging ScaffoldSeq for deep sequencing data analysis, we quantified site-specific amino acid enrichment for each position throughout the CDR-H3 region while simultaneously accounting for varying loop lengths.^47^ In addition, we were able to cluster CDR-H3 sequences from each sorting track into families based on sequence similarity (See *Materials and Methods* for details).

Heatmaps generated using the *seaborn* package in Python depict the degree of enrichment for different amino acids across CDR-H3 (**Figure 3C**). In the b1F1-A-AAZ population, a broader set of amino acids were observed throughout CDR-H3, indicated by relatively small changes in amino acid frequencies relative to the naïve library. For sorting tracks using warheads with moderate or weak potencies, enrichment and depletion of residues within CDR-H3 are more pronounced. For example, tyrosine is enriched at several CDR-H3 positions in multiple populations, especially b1F2-O-BS, B2F2-O-BS, and b1F3-A-AS, and leucine is preferentially enriched at the end of CDR-H3 in the b1F2-O-BS and B2F2-O-BS populations. These results suggest that pharmacophore identity can play a role in shaping CDR-H3 composition. The presence of the potent AAZ pharmacophore appears to be sufficient to enable binding to bCA with a relatively broad set of sequences, whereas the range of sequences compatible with moderate or weak affinity pharmacophores is much narrower by comparison.

In addition to looking at enrichment trends for individual amino acid residues, we also examined the sets of full CDR-H3 sequences identified in each population (**Supplementary Table S3**; see *Materials and methods* for details). Inspection of the ten most frequently identified sequences from each campaign indicates that most CDR-H3 sequences are unique to a specific pharmacophore, except for three sequences shared between alkyl sulfonamide **3** (b1F3-A-AS) and benzene sulfonamide **1** (b1F2-A-BS). Overall, the mostly nonoverlapping top sequences suggests that sorting with distinct pharmacophores leads to differences in individual sequence preferences, not just general amino acid preferences or loop length preferences. We also looked at the total number of unique sequences and total number of sequence clusters identified in each enriched population (**Supplementary Table S3, Supplementary Figure S3**; sequence similarity as defined by an 80% similarity threshold; see *Materials and methods* for details). The population sequenced following sorts with AAZ (b1F1-A-AAZ) contained 2792 unique sequences and 1977 sequence clusters. In contrast, all of the remaining populations were found to contain between 190 and 339 unique sequences and between 24 and 126 clusters. The number of unique sequences is much lower in these populations, and the ratio of clusters compared to unique sequences is much smaller than for the case of AAZ. This data provides further evidence that a much broader range of sequences are compatible with the potent AAZ pharmacophore in comparison to other pharmacophores. Overall, deep sequencing data demonstrates that different pharmacophores lead to distinct preferences in terms of ncAA incorporation site, CDR-H3 loop length, CDR-H3 composition, and specific CDR-H3 sequence. These observations highlight the powerful effects pharmacophores can have on screening outcomes when sorting chemically expanded antibody libraries.

### Yeast surface characterization of exemplary clones

To complement population-wide deep sequencing analyses, we isolated and characterized a series of individual clones (**Supplementary Table S4, Supplementary Figure S4**). Some individual clones were identified multiple times in Sanger sequencing, and in general the extent to which individual clones were repeated is consistent with the deep sequencing data described above (all repeats listed in **Supplementary Table S4**). Flow cytometry analysis revealed that in yeast display format, all pharmacophore-modified individual clones exhibited bCA binding at a concentration of 50 nM, while no bCA binding was detected when the pharmacophore was not installed (**Supplementary Figure S4**). From these data, we identified clones from each population that exhibited high bCA binding levels for further characterization (**Table 1, Figure 4**). Consistent with the high apparent antigen binding, most of these clones were repeated frequently in individual clone analysis and possess attributes (eg., TAG position, CDR-H3 loop length) that were identified in deep sequencing analysis.

**Table 1.**
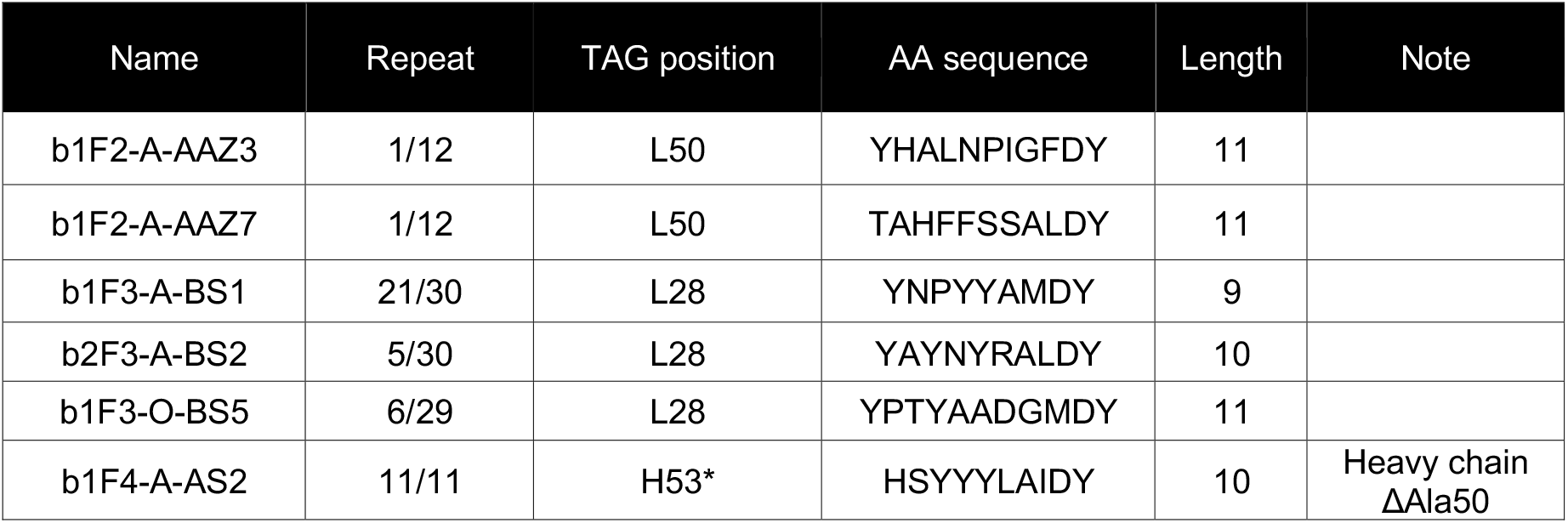
Individual clones characterized in detail in yeast display and biochemical assays.

**Figure 4.**
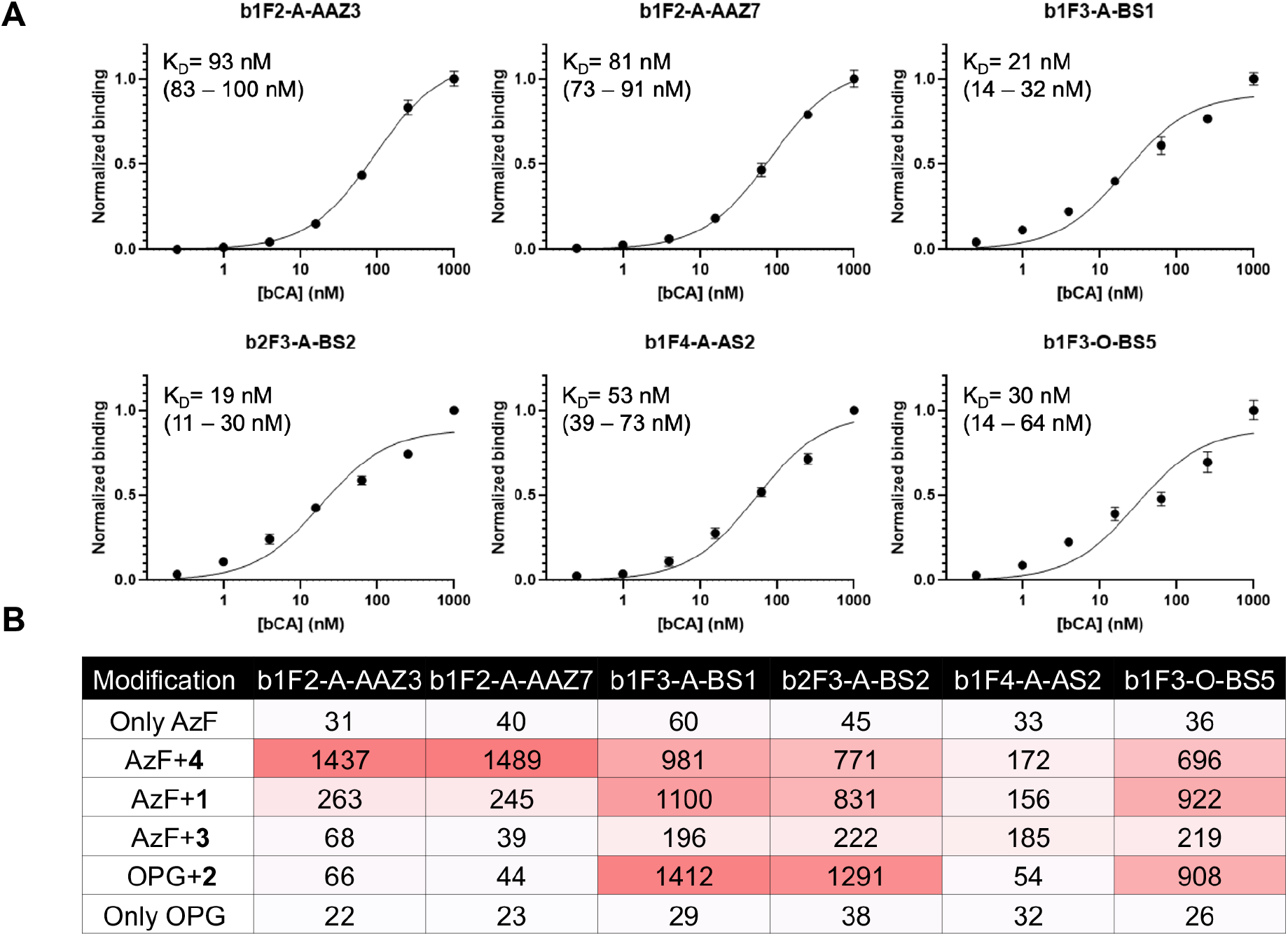
Characterization of clones on the yeast surface. **A)** Yeast surface titrations of clones to determine apparent binding affinities. Numbers in parentheses are the ranges of the 95% confidence intervals determined during analysis of titration data. **B)** “Warhead Swap” binding test of all the clones with different pharmacophores using 25 nM bCA for each displayed sample. Numbers are the median fluorescence intensities of the detected binding bCA (arbitrary units). A warhead swap experiment where cells displaying the same series of hybrids were treated with 250 nM bCA is depicted in the supplementary information (**Supplementary Figure S6**; see also Supplementary Tables S5 and S6 for coefficient of variation data on experiments run at 25 nM and 250 nM bCA).

We performed yeast surface titrations of each hybrid to determine apparent bCA binding constants (**Figure 4A, Supplementary Figure S5**). From these experiments, all apparent K_D_s were determined to be in the double-digit nM affinity range. For the AAZ-containing hybrids, these values are comparable to the potency of the AAZ warhead (K_i_ ∼10 nM).^48^ For the aromatic and aliphatic sulfonamide hybrids, these K_D_ values compare favorably with the reported potencies of the respective pharmacophores (K_i_ ∼100-1000 nM for aromatic benzenesulfonamides and K_i_ ∼ 10,000 nM for the aliphatic sulfonamide).^41-43^ These observations suggest that the antibody scaffolds contribute affinity improvements within isolated hybrids.

To further investigate the importance of pharmacophore-antibody pairings, we prepared hybrids in which the pharmacophores were switched out for each of the other three pharmacophores (along with the corresponding ncAA when necessary) for all 6 of the clones in Table 1. Following this “warhead swap,” we evaluated bCA binding at concentrations of 25 nM bCA (**Figure 4B**; see **Supplementary Table S5** for coefficient of variation data) and 250 nM bCA (**Supplementary Figure S6**; **Supplementary Table S6**). Changes in pharmacophores led to altered binding activity, and in many cases resulted in decreased bCA binding. The AAZ clones (b1F2-A-AAZ3, b1F2-A-AAZ7) show the highest level of binding signal when modified with AAZ, but exhibit dramatically reduced antigen binding levels following conjugation with any of the weaker sulfonamides. This suggests that the binding of these AAZ clones is dependent on the presence of the strong pharmacophore. In contrast, substituting AAZ into clones isolated from the sorts conducted with the less potent pharmacophores results in retention of binding function. Interestingly, AAZ installation in clones isolated from benzenesulfonamide (b1F3-A-BS1, b2F3-A-BS2, and b1F3-O-BS5) or alkyl sulfonamide (b1F4-A-AS2) sorts does not increase the levels of antigen binding detected. In fact, in several cases antigen binding levels are decreased, despite the increased potency of the AAZ pharmacophore compared to the warhead used during screening. In the most pronounced example, we highlight the binding behavior of b1F4-A-AS2: even though the alkyl sulfonamide-modified hybrid exhibits the lowest levels of antigen binding of any individual clone, modifications with the AAZ pharmacophore or aryl sulfonamide warheads did not increase antigen binding levels of this hybrid when treated with 25 nM bCA (At the higher concentration of 250 nM bCA, a maximum of a 2-fold increase in antigen binding level is detected in b1FA-A-AS2 following a warhead swap). These observations provide further evidence for the importance of the warhead-antibody pairing in determining hybrid binding activity.

We have previously observed that the substitution of AzF + **1** with OPG + **2** in fibronectin-based hybrids can lead to a loss of binding in human isoform-specific clones^32^. However, in the case of the bCA-binding hybrids isolated in this work, the AzF + benzenesulfonamide (b1F3-A-BS1, b2F3-A-BS2) and OPG + benzenesulfonamide (b1F3-O-BS5) clones exhibit high levels of bCA binding when conjugated with either **1** or **2** (with AzF or OPG, respectively). Apparently, presumed changes in the polarity of the triazole linkage as well as the presence (or absence) of the extra oxygen atom in the linker does not have substantial effects on binding. This could be due to the fact that 1) **1** and **2** are both benzenesulfonamides; and 2) the three clones all have the same TAG codon position, meaning the pharmacophores are presented at the same antibody location within each of the three hybrids. Finally, the warhead swap data again confirms that no bCA binding is detected in the absence of a pharmacophore modification. Overall, these data show that isolated clones usually exhibit the highest binding levels when modified with the pharmacophore used during sorting. This provides further evidence for pharmacophore-dependent hybrid binding and highlights the importance of the antibody scaffold in determining hybrid performance.

### Solution-based characterization of exemplary clones

We prepared soluble forms of hybrids to investigate inhibitory properties in biochemical assays. ScFv-Fc forms were cloned and then produced with a previously described yeast secretion system that supports production of proteins with ncAAs (**Figure 5A**).^49^ Following expression and subsequent Protein A purification, we verified construct isolation using SDS-PAGE. Consistent with our prior work, coomassie-stained gels show banding patterns indicating a portion of sample running at approximately 70 kDa, and a portion running at a higher molecular weight that corresponds to a glycosylated form of the same product^35, 39, 44, 49, 50^.

**Figure 5.**
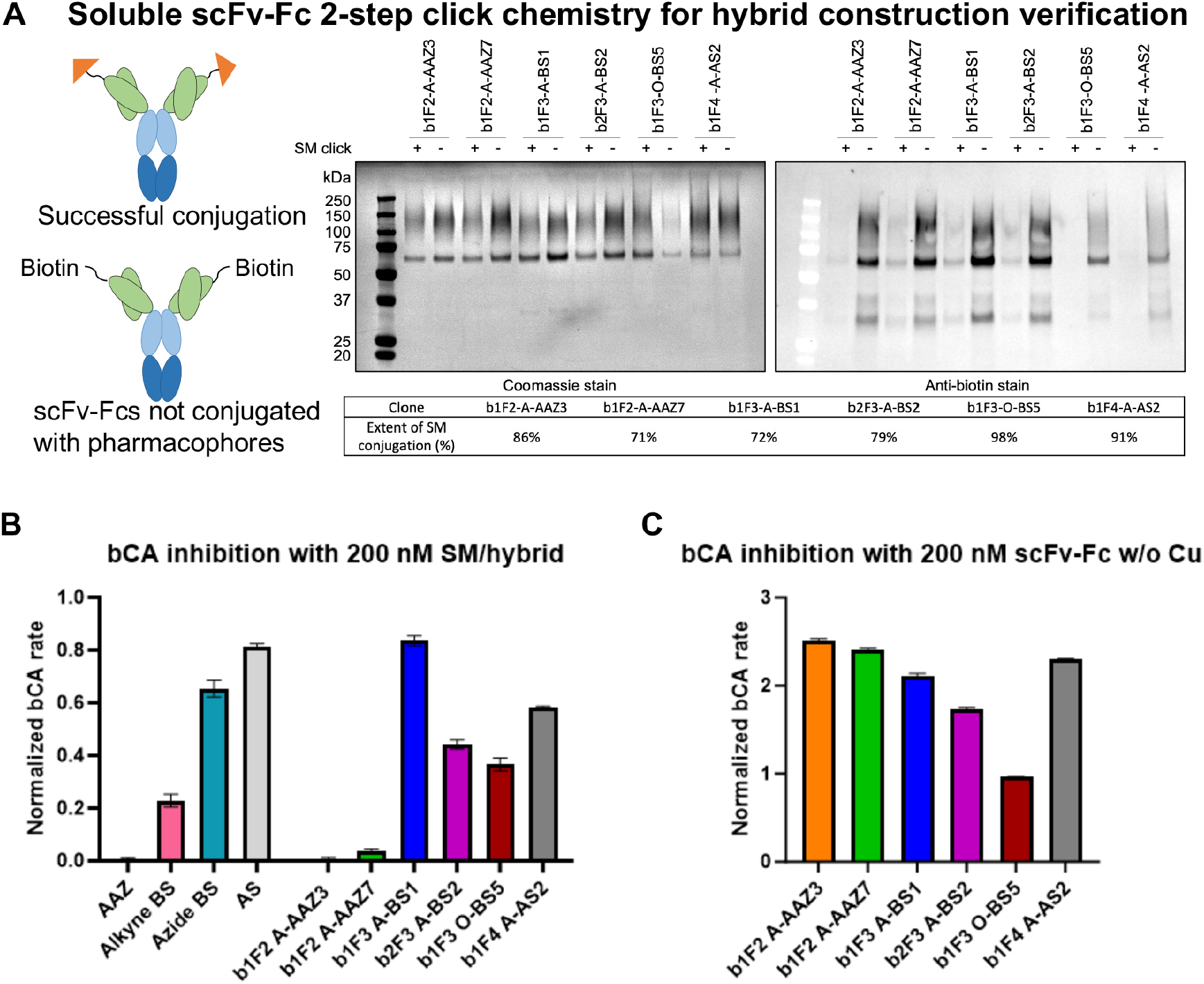
Soluble hybrid construction and characterization. **A)** 2-step click chemistry evaluation of hybrid construction in solution. SM: small molecule. Extents of reaction were calculated using ImageJ (see **Supplementary Figure S7** for images and *Materials and methods* for details). **B)** Single-point inhibition assay with 200 nM small molecule (SM) or hybrid in bCA inhibition assays. bCA concentration is 50 nM, and the substrate is 2 mM 4-nitrophenyl acetate. Experiments were performed in technical triplicates. **C)** Single-point inhibition assay with 200 nM scFv-Fc treated with reaction mixtures while omitting copper (Cu). Reaction rate is normalized to the rate of 50 nM bCA with 2 mM 4-NPA in buffer.

We installed pharmacophores within purified scFv-Fcs using CuAAC and estimated the efficiency of installation using our previously described 2-step click chemistry approach (see the *Materials and methods* section *ScFv-Fc click chemistry* for details).^39^ SDS-PAGE and Western blot experiments indicated the installation of pharmacophores with a range of 71% to 98% efficiency (**Figure 5A, Supplementary Figure S7**). These data strongly suggest that the desired hybrids were successfully constructed in soluble form.

We then investigated whether the soluble hybrids inhibit bCA enzymatic activity (**Figure 5B, Supplementary Figure S8, S9, Supplementary Table S9**). Using 4-nitrophenyl acetate as a colorimetric reporter of bCA activity, we conducted activity assays in the presence of hybrid inhibitors. In general, the results show that all 6 hybrids show pharmacophore-dependent inhibition of bCA in solution. At 200 nM hybrid, we observe evidence for the inhibition of bCA (**Figure 5B**) for all six soluble hybrids evaluated. Notably, clones b1F3-O-BS5 and b1F4-A-AS2 (which contain weak sulfonamide pharmacophores **2** and **3**, respectively) exhibit increased target inhibition as hybrids in comparison to controls containing pharmacophores alone at 200 nM. These are noteworthy findings since only binding was assayed during screening and individual clone validation. Concentration-response plots for hybrids indicate varied behaviors (**Supplementary Figure S8**). We observe monotonic decreases in bCA activity as AAZ-modified hybrid concentration increases, but bCA activity varies in more complex ways as hybrids modified with weaker pharmacophores are added at increasing concentrations. In several cases, bCA activity increases at mid-range concentrations (50 nM and 100 nM) before decreasing again at 200 nM. In additional control experiments, samples in which bCA was treated with unconjugated scFv-Fcs at 200 nM have elevated activity in comparison to untreated controls (**Figure 5C, Supplementary Figure S9**). These unconjugated scFv-Fcs were subjected to the same procedures used to produce the conjugated hybrids, except for omission of the copper catalyst used in CuAAC reaction (CuSO_4_ and chelating ligands, **Supplementary Table S9**). Similarly, scFv-Fcs that have not been treated with any CuAAC reagents enhance bCA activity as well (**Supplementary Figure S9**). These observations suggest that the proteins on their own may enhance bCA activity, providing a potential explanation for the nonmonotonic changes in enzyme activity observed in the concentration-response plots. However, further work will be needed to gain mechanistic understanding of these phenomena. Overall, biochemical assays revealed that discovered hybrids can exhibit inhibitory properties, even when presenting moderate or weak potency pharmacophores within antibody structures. In addition, enzymatic assays revealed that antibody components of hybrids do not exhibit detectable enzyme inhibition in the absence of pharmacophores. Thus, sequencing data, on-yeast characterizations, and in-solution characterizations all demonstrate that installing pharmacophores within antibodies strongly shapes hybrid discovery campaigns.

## Discussion

In this work, we constructed multiple hybrid forms of a billion-member, chemically expanded yeast display library and screened the resulting hybrid collections. Enrichments with pharmacophore-modified antibody libraries led to the isolation of a wide range of clones binding to bCA, with distinct pharmacophores appearing to drive the discovery process into different regions of sequence space. The potent pharmacophore AAZ supported hybrid discovery with diverse collections of antibodies and ncAA attachment points. Weaker pharmacophores also supported hybrid discovery, but with narrower sets of preferred pharmacophore modification sites, antibody CDR-H3 loop lengths, and CDR-H3 sequences. We observed little to no sequence overlap between the frequently isolated sets of clones from different pharmacophore tracks, providing further evidence for pharmacophore-driven preferences at the population level. Characterization of individual clones revealed apparent binding constants in the double-digit nanomolar range, even when using moderate potency (benzenesulfonamide) or weak potency (alkyl sulfonamide) pharmacophores. Warhead swap experiments indicated that clones isolated in high potency warhead sorting tracks exhibit diminished binding capacity upon substitution with less potent warheads; clones isolated from moderate or weak potency warhead tracks generally retain binding function upon swapping. For all hybrids evaluated in solution, we observed inhibition of enzyme activity, and in some cases our data suggests improved inhibitory properties compared to the properties of the warhead alone. Overall, many lines of evidence indicate that distinct pharmacophores substantially shape the discovery efforts reported here, and provide numerous opportunities for further extensions.

It is noteworthy that we were able to isolate hybrids starting from moderate to weak potency pharmacophores. Benzenesulfonamide and alkyl sulfonamide pharmacophores exhibit potencies similar to those of initial hits in many small molecule screens. Our successes in improving the properties of these hits raise the possibility that well-characterized fragments, ligands, and probes from numerous prior efforts may serve as strong “chemical starting points” for initiating pharmacophore-driven antibody discovery^33, 51-61^.

The findings in this manuscript also have implications for binding protein diversities for use in future hybrid library designs. In the antibody diversification scheme used here^35^, the pharmacophore is positioned outside of the diversified binding region, in contrast to previous hybrid libraries where pharmacophores are integrated within diversified binding regions of proteins^30, 31^ or peptides^62-68^. Our successes in isolating hybrids with pharmacophores distal to diversified loops indicates that there are many possible ways to present pharmacophores alongside diversified protein paratopes while also being conceptually consistent with prior work. Most notably, screens of fibronectin-based hybrid libraries revealed CA isoform-dependent preferences in the location of the AAZ pharmacophore attachment site.^31^ While there are many differences between our study and these previous studies (including antibody vs fibronectin, three pharmacophores vs one, pharmacophore distal from paratope vs integrated, bCA versus isoforms of human CA targets), all of these findings indicate the importance of carefully considering the positioning of pharmacophores within future hybrid libraries to maximize the potential for the discovery of potent, function-disrupting hybrids.

Pharmacophore-driven antibody discovery is a promising addition to antibody engineering and hybrid screening approaches. Even with the relatively modest performance of the simple antibody diversity used in this study, the introduction of pharmacophores facilitated discovery of a broad set of hybrids. Efforts to build second-generation antibody hybrid libraries that incorporate additional features of high-performing antibody libraries are underway in our laboratory.^69-75^ In addition, the flexibility of yeast display can be used to extend hybrid diversity beyond antibodies and fibronectins^30, 31^ to include protein binding scaffolds such as nanobodies^76-80^, affibodies^81^, and Sso7d^82^; complex peptide display formats have also recently been validated in yeast display format^83^. The availability of diverse formats will enable broader investigations of the interplay between pharmacophore and polypeptide diversity that are needed to elucidate fundamental principles of hybrid design and screening. Finally, hybrid discovery and characterization approaches can be used to generate sequencing and structural data needed to apply advances in biomolecular machine learning^84-86^ and computational protein design^87-90^ to unexplored hybrid sequence spaces. Our demonstration of pharmacophore-driven antibody discovery highlights opportunities to pursue hybrids as research tools and potential therapeutic leads for poorly drugged targets.

## Acknowledgments

This work was supported by a grant from the National Institute of General Medical Sciences of the National Institutes of Health (R35GM133471 to J.A.V.), the Bright Futures Assistant Professorship at Tufts University (to J.A.V.), a grant from the National Institutes of Health (DP2HD91798 to N.U.N.), a grant from the National Science Foundation (#1935354 to N.U.N.), and Tufts Launchpad | Accelerator (to N.U.N.). The content is solely the responsibility of the authors and does not necessarily represent the official views of the National Institutes of Health or Tufts University.

