## Supporting information for "Pharmacophore-driven antibody discovery on the yeast surface"

#### Supplementary Figures

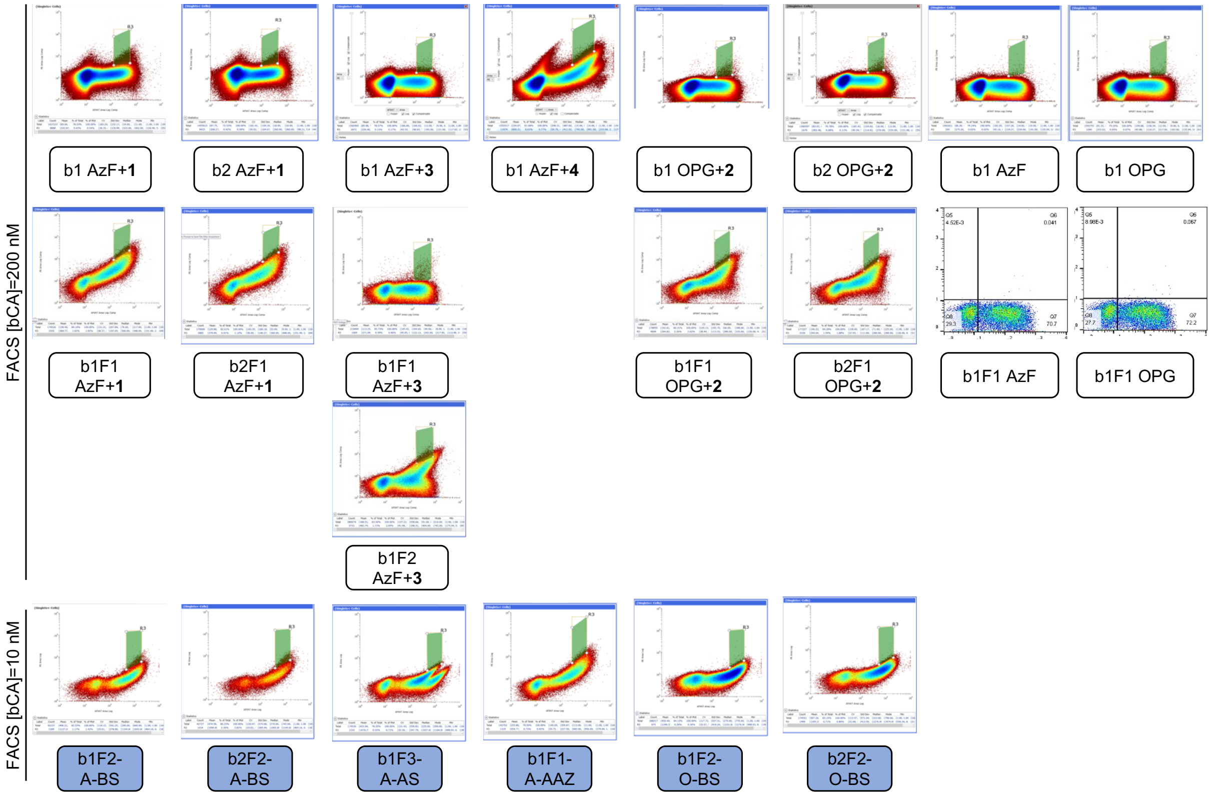


**Supplementary Figure S1.** Populations sorted and gates used during FACS. Gates were set to sort for approximately the top 0.5% of binders (high detection of anti-biotin PE/biotinylated bCA on the y axis), avoiding low read through clones (low cMyc/AF647 detection on the x axis) and potential doublets (high cMyc/AF647 detection on the x axis).

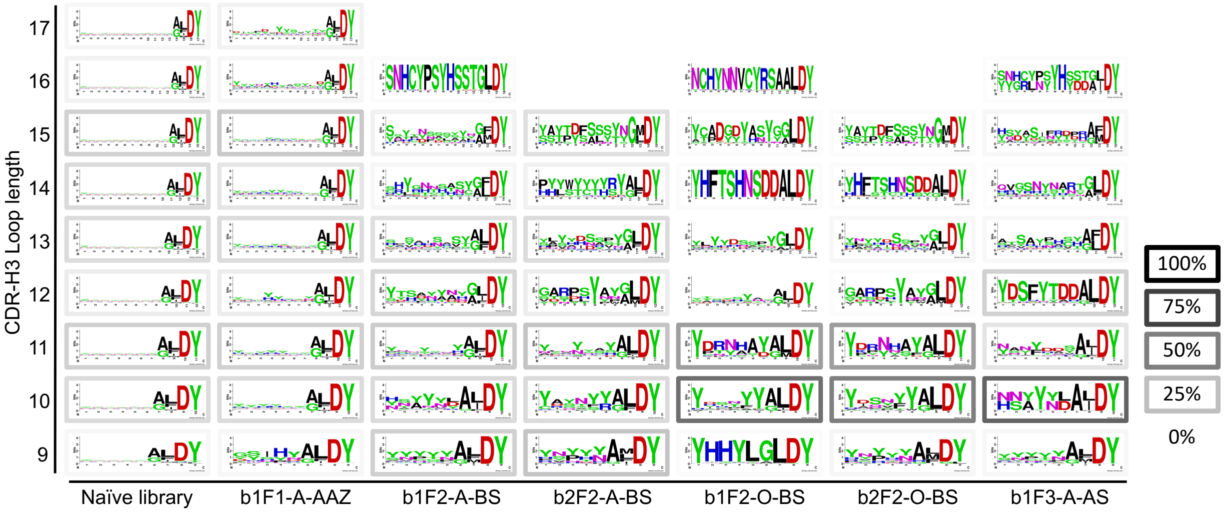


**Supplementary Figure S2.** Sequence logos of the different populations as a function of length. Outlines surrounding the individual logos reflect the percentage of the specific length in the sequenced population.

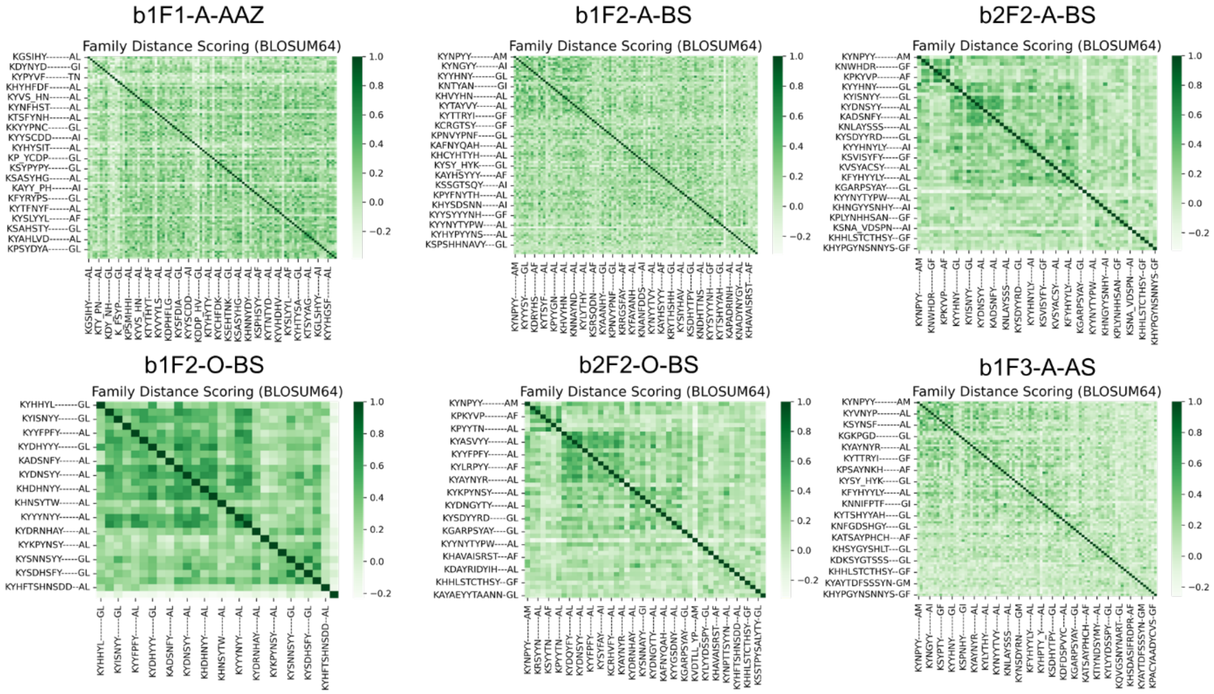


**Supplementary Figure S3.** Family distance scoring matrices (BLOSUM64 scores) for the different sorting populations. Each heatmap (generated in Python using the *seaborn* package) compares the sequence similarity of top clones from each cluster. A higher score (darker green) corresponds to more closely related sequences while a lower score (lighter green) corresponds to two sequences that are more distantly related.

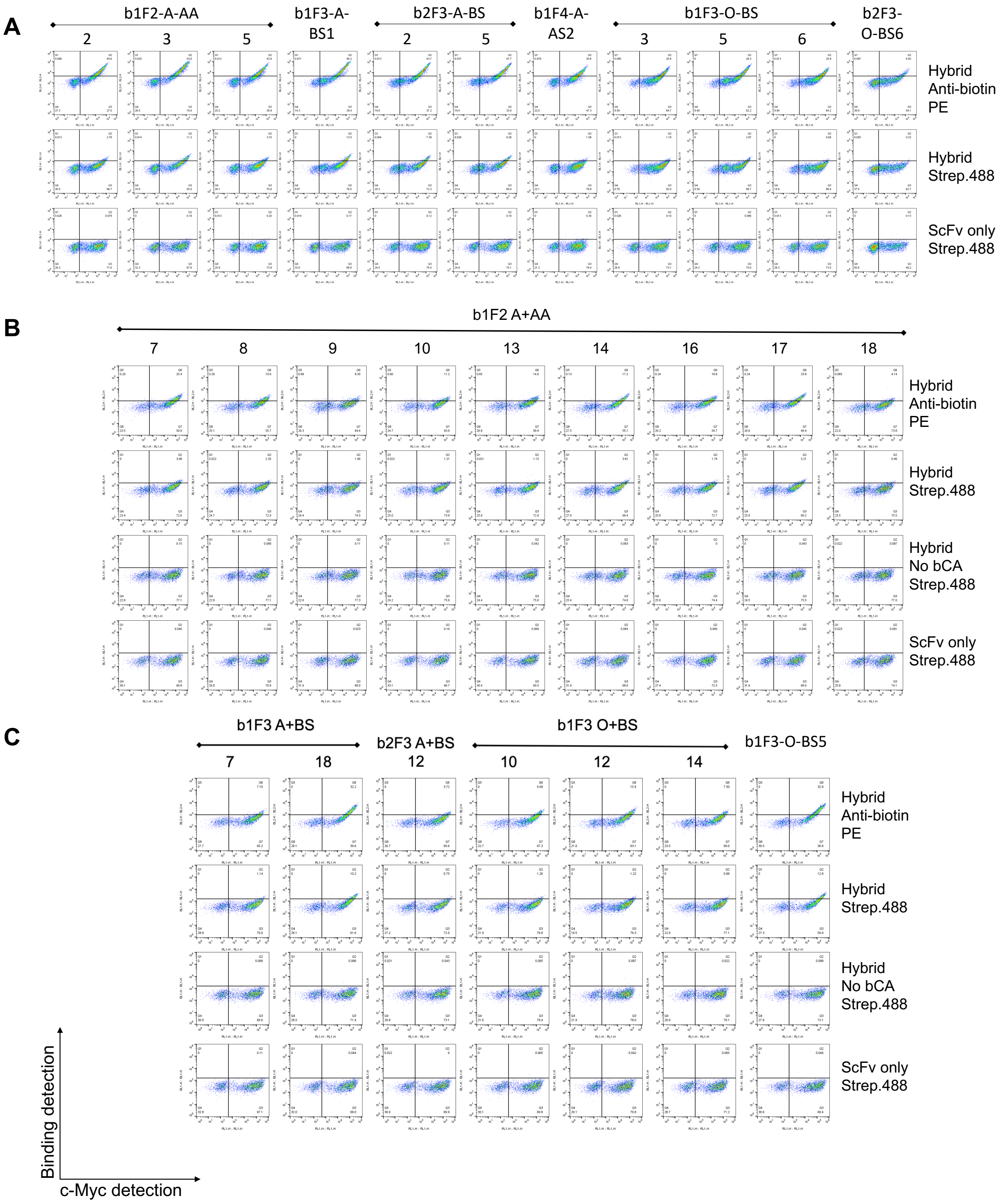


**Supplementary Figure S4.** Binding assays of individual clones treated with 50 nM bCA. Hybrid: scFvs conjugated to corresponding pharmacophore used during sorting. The data presented in (**A**) were collected on different days from the data presented in (**B**) and (**C**). Anti-biotin PE: mouse anti-biotin antibody conjugated to PE (used during cell sorting). Strep.488: Streptavidin conjugated to Alexa Fluor 488.

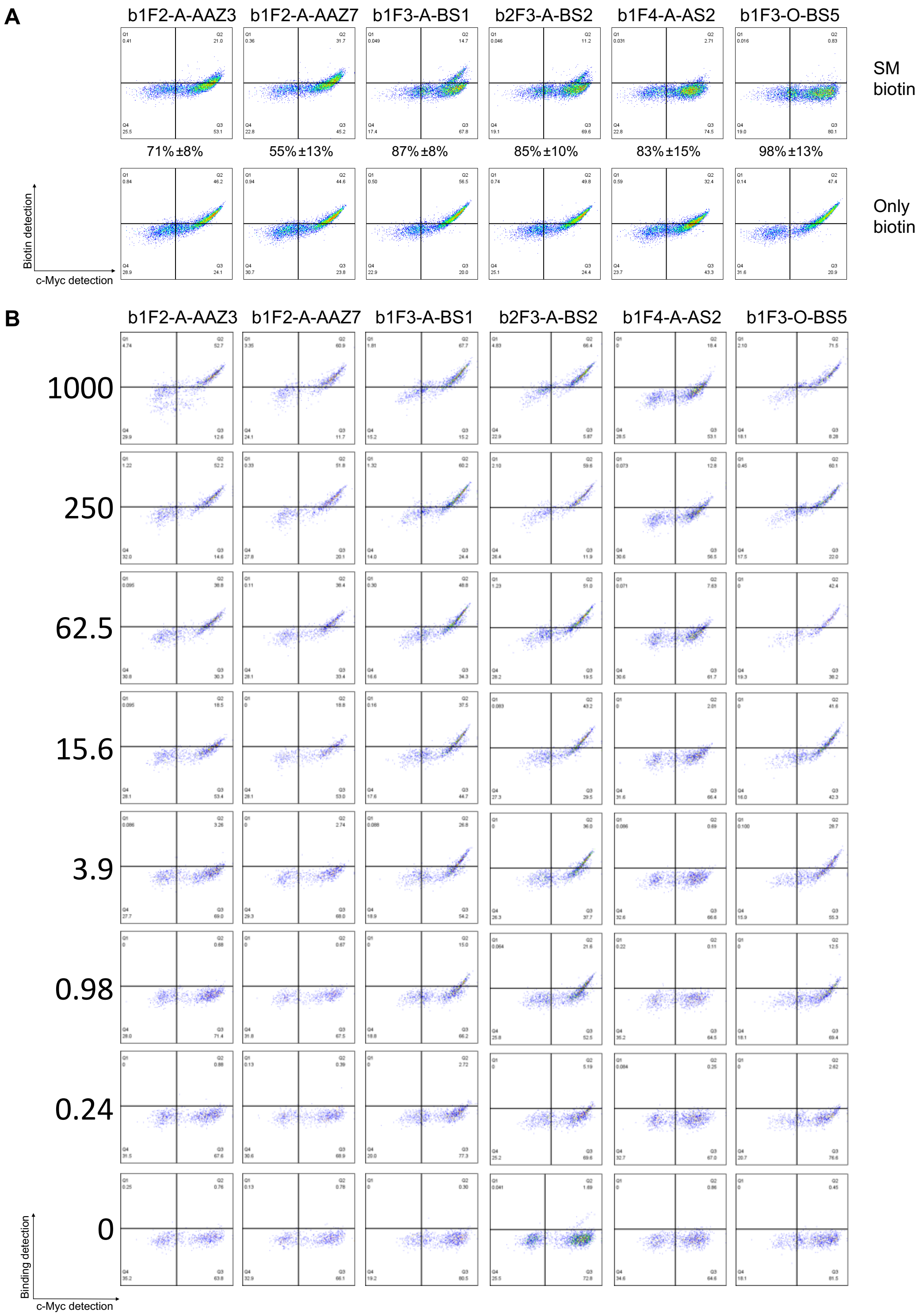


**Supplementary Figure S5.** Yeast surface titrations of exemplary clones. **A**) 2-step click chemistry to evaluate efficiency of hybrid construction on yeast surface. The extent of reaction with propagated error is listed below each of the respective flow cytometry dot plots. **B**) Dot plots for yeast surface titrations. Experiment was done in triplicates; only one replicate is shown here.

**A**

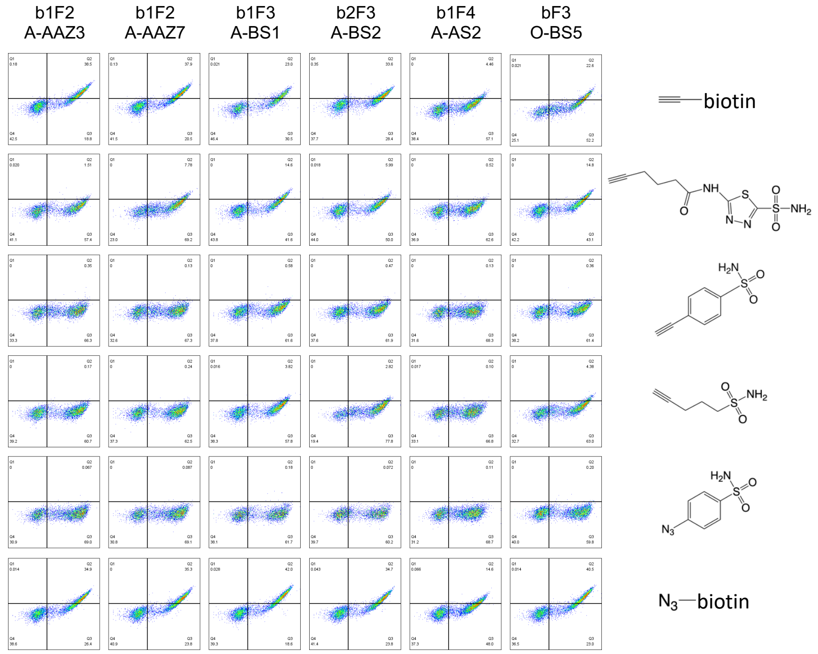


**B**

| Modification | b1F2-A-AAZ3 | b1F2-A-AAZ7 | b1F3-A-BS1 | b2F3-A-BS2 | b1F4-A-AS2 | b2F3-O-BS5 |
|---|---|---|---|---|---|---|
| AzF+**4** | 91%±7% | 78%±7% | 35%±14% | 65%±11% | 63%±37% | 21%±21% |
| AzF+**1** | 97%±7% | 97%±6% | 85%±12% | 91%±10% | 89%±36% | 87%±15% |
| AzF+**3** | 96%±6% | 97%±6% | 71%±12% | 75%±11% | 81%±35% | 63%±17% |
| OPG+**2** | 98%±7% | 98%±7% | 99%±6% | 99%±7% | 95%±18% | 99%±5% |

**C**

| Modification | b1F2-A-AAZ3 | b1F2-A-AAZ7 | b1F3-A-BS1 | b2F3-A-BS2 | b1F4-A-AS2 | b1F3-O-BS5 |
|---|---|---|---|---|---|---|
| Only AzF | 28 | 46 | 122 | 62 | 19 | 37 |
| AzF+**4** | 1635 | 1607 | 808 | 805 | 439 | 1135 |
| AzF+**1** | 642 | 595 | 1533 | 1258 | 496 | 1510 |
| AzF+**3** | 150 | 55 | 402 | 371 | 198 | 371 |
| OPG+**2** | 288 | 229 | 1431 | 1315 | 171 | 1189 |
| Only OPG | 24 | 27 | 79 | 67 | 47 | 50 |

**Supplementary Figure S6.** Pharmacophore swap of exemplary clones. **A)** 2-step click chemistry verification of hybrid construction on yeast surface. **B)** Extent of reaction calculated from (**A**). **C)** Pharmacophore swap of exemplary clones labeled at 250 nM bCA; see main text **Figure 4B** for clones labeled at 25 nM bCA, and Supplementary Tables S5 and S6 for coefficient of variant data for all samples.

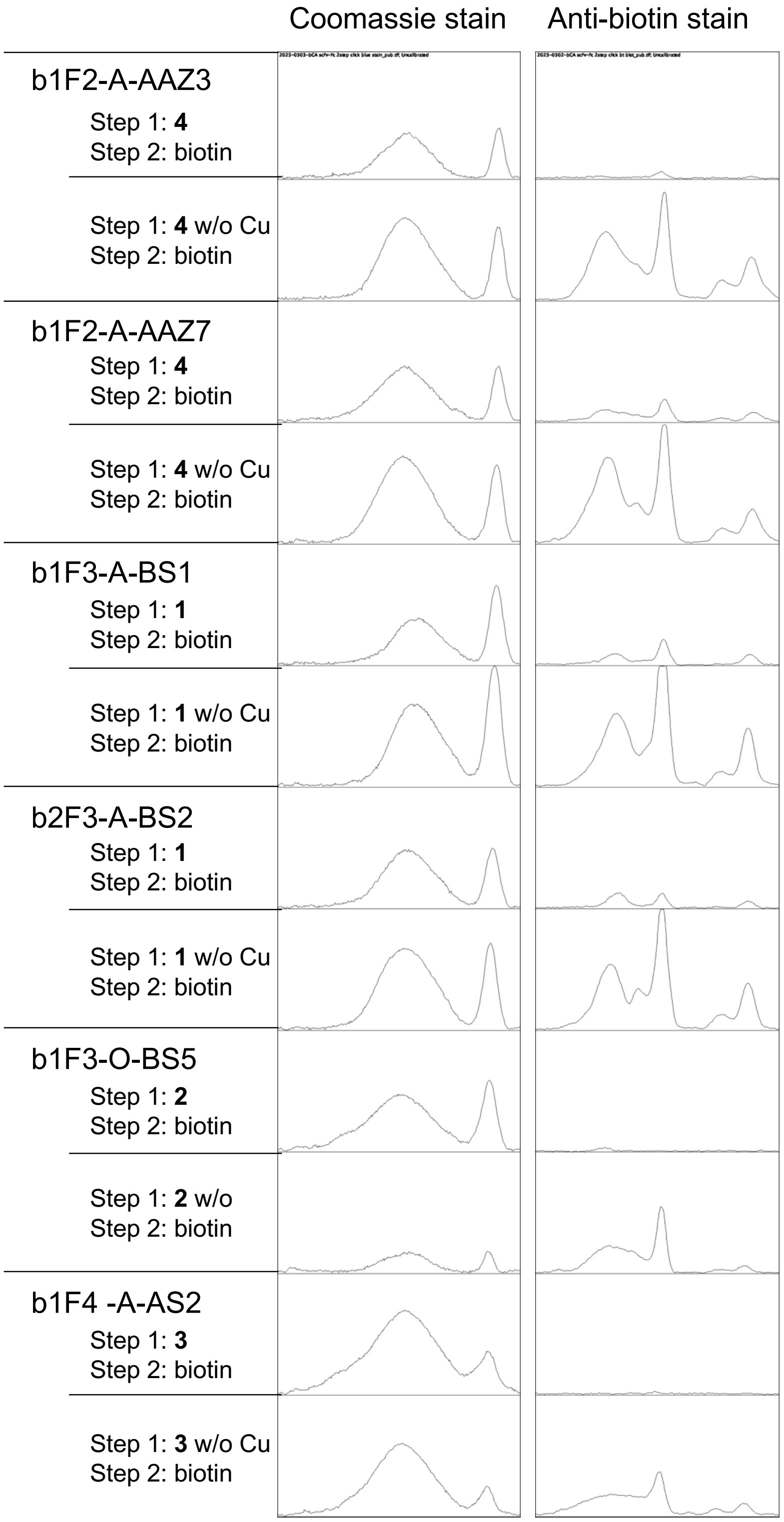


**Supplementary Figure S7.** Peaks from images of the western blot and SDS-PAGE shown in Figure 5A. The images were analyzed with Image J. The background was subtracted using a rolling ball radius of 50 pixels and light background, and the lanes were assigned manually and plotted with the gel analysis tool.

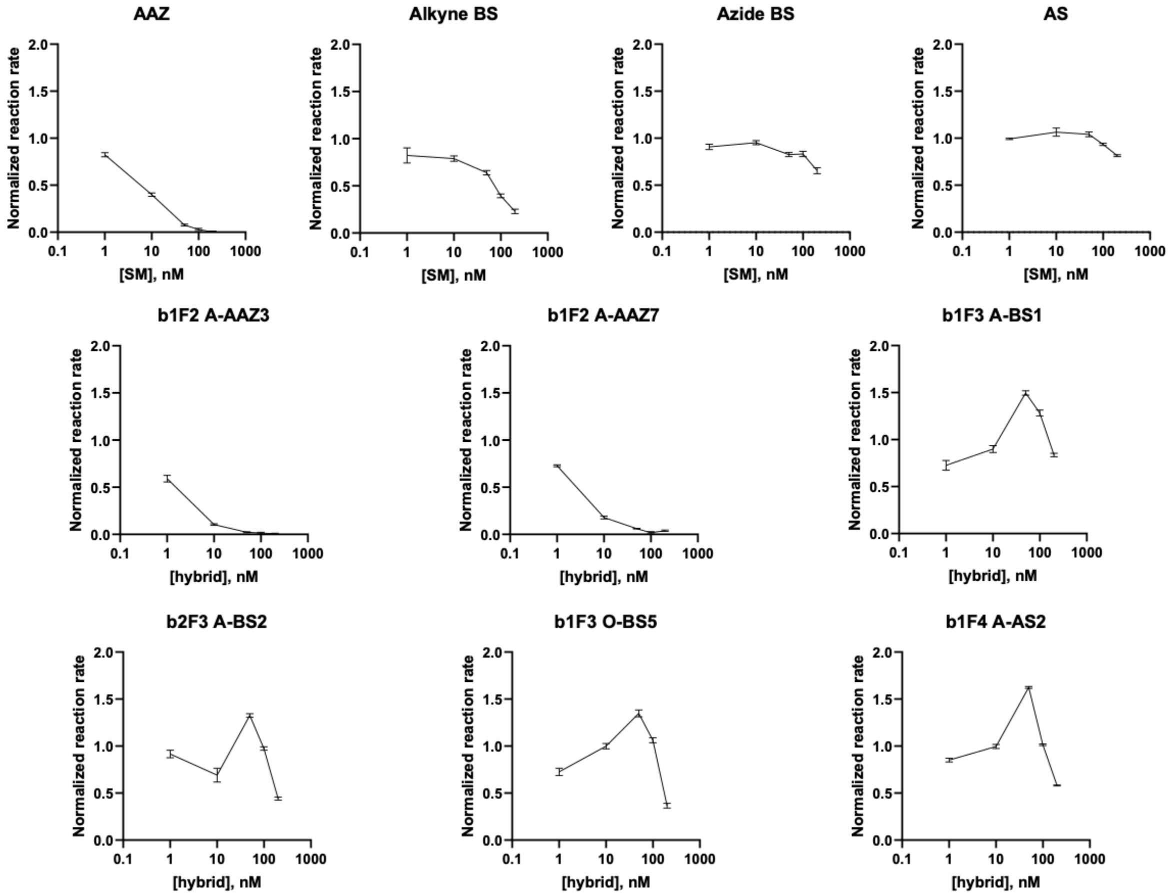


**Supplementary Figure S8.** Concentration-response curves of pharmacophores and hybrids at hybrid concentrations of 0 nM, 1 nM, 10 nM, 50 nM, 100 nM, 200 nM. [bCA] = 50 nM, [4-NPA] = 2 mM.

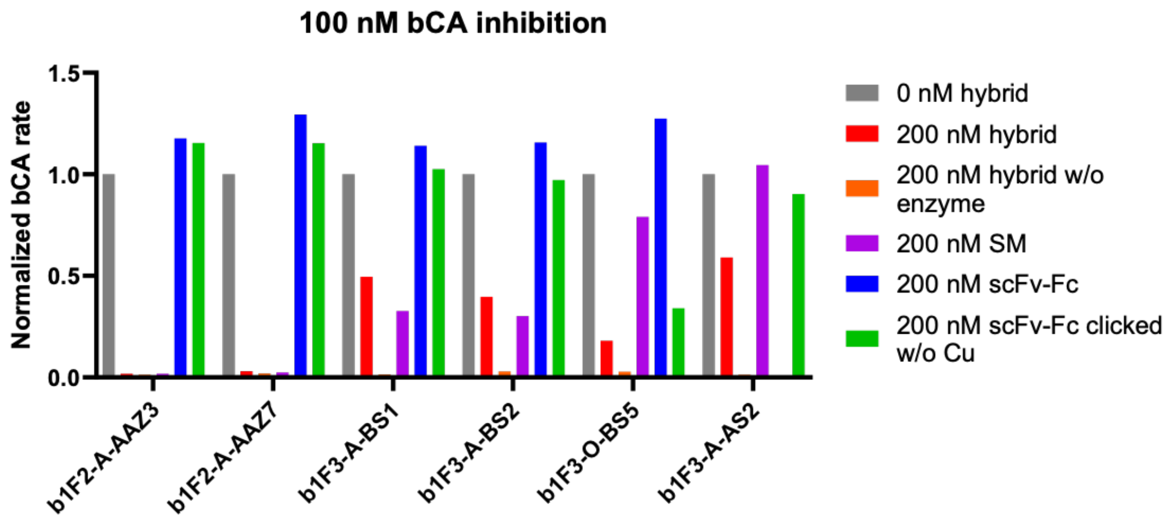


**Supplementary Figure S9.** Single concentration inhibition assay. Detailed descriptions of each condition are listed in **Supplementary Table S7**, with the exception of the bCA concentration, which was 100 nM in this series of assays. For b1F3-A-AS2, the 200 nM scFv-Fc sample was not tested due to limited amount of sample at the time of experiment.

### Supplementary Tables

**Supplementary Table S1.** DNA sequences of the scFv frameworks used for synthetic antibody library construction and human Fc used for secretion constructs. The NheI and BamHI restriction sites introduced before and after the scFv frameworks to allow cloning into a pCTCON2 vector are highlighted. The diversified CDR-H3 sequence is in the heavy chain after the IGHV3-23 and before the JH4 domain.

| Domain | Sequence |
|---|---|
| VL: IGKV1-39-JK4 | GCTAGCGACATACAGATGACTCAAAGTCCCAGTTCACTATCTGCGTCTGTTGG<br>TGATAGAGTCACCATTACGTGTAGAGCTTCTCAGTCGATTAGCTCGTACTTGAA<br>TTGGTATCAACAGAAACCAGGGAAAGCTCCAAAGTTGCTGATCTATGCAGCAT<br>CTAGCTTACAAAGTGGTGTACCTTCCAGGTTTTCAGGCTCAGGATCTGGAACT<br>GATTTCACACTTACCATATCATCCTTACAACCGGAAGATTTCGCCACATATTAC<br>TGCCAACAATCCTACTCTACTCCACCTACATTTGGTGGTGGCACTAAAGTGGA<br>GATTAAGGGTACTACTGCCGCTAGTGGTAGTAGTGGTGGCAGTAGCAGTGGT<br>GCC |
| SidLink (between VL and VH) | GGTACTACTGCCGCTAGTGGTAGTAGTGGTGGCAGTAGCAGTGGTGCC |
| VH: IGHV3-23 | GGTACTACTGCCGCTAGTGGTAGTAGTGGTGGCAGTAGCAGTGGTGCCGAGG<br>TGCAATTGCTAGAATCAGGAGGTGGTTTGGTACAACCTGGTGGTAGCTTAAGG<br>TTGTCTTGTGCTGCTAGTGGATTCACGTTTAGTAGCTATGCCATGTCATGGGTT<br>AGACAAGCTCCAGGTAAAGGCTTAGAATGGGTTTCTGCGATATCTGGATCTGG<br>TGGGTCAACTTACTATGCAGATTCCGTCAAAGGCAGATTTACCATTTCCAGAGA<br>CAATTCGAAGAATACACTGTACCTTCAGATGAACTCGTTACGTGCAGAAGATAC<br>TGCTGTTTACTACTGTGCTAAG |
| VH: JH4 | GACTACTGGGGCCAAGGAACCCTGGTCACCGTCTCCTCAGGATCC |
| Human Fc | GCTCCAGAATTGTTAGGTGGTCCCAGCGTCTTTTTGTTCCCGCCAAAGCCGAA<br>AGACACGTTGATGATCAGTAGAACCCCGGAAGTAACATGTGTTGTTGTGGACG<br>TGAGTCACGAGGACCCCGAAGTAAAGTTTAATTGGTACGTGGATGGTGTTGAA<br>GTTCACAACGCCAAGACTAAGCCTAGAGAAGAACAGTACAATTCAACATATCG<br>TGTTGTTAGCGTGTTGACTGTTCTTCACCAAGATTGGCTTAATGGTAAGGAATA<br>CAAATGCAAAGTATCCAATAAAGCACTTCCAGCTCCGATAGAGAAGACAATCA<br>GTAAAGCCAAAGGTCAGCCCAGAGAACCTCAAGTTTATACCTTGCCTCCGTCT<br>AGAGACGAATTGACCAAAAATCAAGTCTCCCTGACTTGCCTAGTGAAGGGCTT<br>TTACCCATCCGATATAGCGGTAGAATGGGAATCGAATGGACAACCGGAAAACA<br>ACTATAAGACTACTCCCCCTGTATTGGATTCTGATGGATCTTTCTTTTTGTATAG<br>CAAATTGACCGTTGATAAGTCTAGGTGGCAGCAGGGAAACGTCTTCTCATGTA<br>GTGTTATGCACGAAGCCTTACACAATCACTACACACAAAAGTCATTATCCTTGT<br>CTCCAGGTAAGAGATCT |

**Supplementary Table S2.** FACS population click chemistry extents of reaction determined by two-step click chemistry experiments. Due to the high level of noise in samples acquired on the s3e sorter, the resulting CV values are larger than the median fluorescence intensity; this prevents calculation of propagated error.

| Population | Data collection machine | Extent of click reaction | Propagated error |
|---|---|---|---|
| b1-A-BS | S3e cell sorter | 83% | |
| b2-A-BS | | 86% | |
| b1-A-AS | | 74% | |
| b1-A-AAZ | | 39% | |
| b1-O-BS | | 96% | |
| b2-O-BS | | 97% | |
| b1F1-A-BS | | 91% | |
| b2F1-A-BS | | 90% | |
| b1F1-A-AS | | 86% | |
| b1F1-A-AAZ | | 68% | |
| b1F1-O-BS | | 94% | |
| b2F1-O-BS | | 94% | |
| b1F2-A-AS | | 88% | |
| b1F2-A-BS | Attune flow cytometer | 87% | 19% |
| b2F2-A-BS | | 88% | 11% |
| b1F3-A-AS | | 91% | 11% |
| b1F2-O-BS | | 93% | 11% |
| b2F2-O-BS | | 96% | 9% |

**Supplementary Table S3.** Highest frequency sequences identified in ScaffoldSeq analysis.

| **b1F1-A-AAZ** | | | **b1F2-A-BS** | | | **b2F2-A-BS** | | |
|---|---|---|---|---|---|---|---|---|
| *2792 unique proteins* | *1977 clusters* | | *339 unique proteins* | *126 clusters* | | *236 unique proteins* | *52 clusters* | |
| **CDR-H3 Sequence** | **Frequency** | **Additional Top-10 Occurances** | **CDR-H3 Sequence** | **Frequency** | **Additional Top-10 Occurances** | **CDR-H3 Sequence** | **Frequency** | **Additional Top-10 Occurances** |
| KHFFYYKK-----GL | 760 | 0 | KHSYYYL------AI | 1090 | 1 (b1F3-A-AS) | KYNPYY-------AM | 1612 | 1(b1F2-A-BS) |
| KYYHFHDPNY---GL | 451 | 0 | KNNAYND------AL | 1015 | 1 (b1F3-A-AS) | KYAYTDFSSSYN-GM | 1512 | 0 |
| KSHHSYYAYGH--AF | 204 | 0 | KYTHANAAY----GI | 1006 | 0 | KYAYNYR------AL | 947 | 0 |
| KSHSTD_TYAYT-AI | 115 | 0 | KYNPYY-------AM | 743 | 1 (b2F2-A-BS) | KSSTPYSALYTY-GL | 873 | 0 |
| KAYHTYAS-----AI | 96 | 0 | KSPSHHNAVY---GL | 742 | 1 (b2F2-A-BS) | KNLAYSSS-----AL | 855 | 0 |
| KSNHHYSHSYNA-GL | 95 | 0 | KSTSDHYNN----AL | 624 | 0 | KPYYWYYYYRY--AL | 748 | 0 |
| KHYHFDF------AL | 92 | 0 | KYYSYYYNH----GF | 595 | 0 | KYLYYDSSPY---GL | 736 | 0 |
| KYDYLCNYKRDD-AL | 87 | 0 | KSYYTN-------AF | 589 | 0 | KGARPSYAY----GL | 729 | 0 |
| KRAFDDHYQSRV-GI | 86 | 0 | KHAVAISRST---AF | 584 | 0 | KHVYHN-------AL | 703 | 0 |
| KYHEYHAY-----AF | 85 | 0 | KYDSFYTDD----AL | 491 | 1 (b1F3-A-AS) | KSPSHHNAVY---GL | 626 | 1 (b1F2-A-BS) |
| **b1F2-O-BS** | | | **b2F2-O-BS** | | | **b1F3-A-AS** | | |
| *190 unique proteins* | *24 clusters* | | *228 unique proteins* | *40 clusters* | | *297 unique proteins* | *80 clusters* | |
| **CDR-H3 Sequence** | **Frequency** | **Additional Top-10 Occurances** | **CDR-H3 Sequence** | **Frequency** | **Additional Top-10 Occurances** | **CDR-H3 Sequence** | **Frequency** | **Additional Top-10 Occurances** |
| KYDRNHAY-----AL | 4603 | 1 (b2F2-O-BS) | KYDRNHAY-----AL | 5995 | 1 (b1F2-O-BS) | KNNAYND------AL | 7367 | 1 (b1F2-A-BS) |
| KYHITYY------AL | 1925 | 0 | KYISNYY------GL | 4424 | 1 (b1F2-O-BS) | KHSYYYL------AI | 7245 | 1 (b1F2-A-BS) |
| KYISNYY------GL | 1866 | 1 (b2F2-O-BS) | KYDQYFY------AL | 3186 | 0 | KYDSFYTDD----AL | 3546 | 1 (b1F2-A-BS) |
| KYPTYAAD-----GM | 1672 | 0 | KYKPYNSY-----AL | 3125 | 0 | KNANFDDS-----AI | 1142 | 0 |
| KYSYFAY------AI | 1015 | 0 | KYASVYY------AL | 2667 | 1 (b1F2-O-BS) | KYSIYHAV-----AL | 400 | 0 |
| KYYFPFY------AL | 995 | 1 (b2F2-O-BS) | KYDNSYY------AL | 1944 | 0 | KCNRSDL------AF | 352 | 0 |
| KYASVYY------AL | 966 | 1 (b2F2-O-BS) | KHNSYTW------AL | 1228 | 0 | KYTSHYYAH----GL | 294 | 0 |
| KYDHYYY------GL | 822 | 0 | KYYFPFY------AL | 1222 | 1 (b1F2-O-BS) | KYNYYTVY-----AL | 273 | 0 |
| KYLRPYY------AF | 682 | 0 | KADSNFY------AL | 913 | 1 (b1F2-O-BS) | KATSAYPHCH---AF | 259 | 0 |
| KADSNFY------AL | 612 | 1 (b2F2-O-BS) | KYSNNAYY-----GI | 793 | 0 | KHSDASIFRDPR-AF | 218 | 0 |

**Supplementary Table S4.** Sequences of all individual clones isolated from the library.

| Sequence name | Repeat | TAG position | H3 Loop sequence | AA sequence | Length | Note |
|---|---|---|---|---|---|---|
| b1F2-A-AAZ 2 | 1/12 | H31 | TACTCCCTTTATTACCTCGCTTTCGACTAC | YSLYYLAFDY | 10 | |
| b1F2-A-AAZ 3 | 1/12 | L50 | TACCATGCTCTCAACCCGATCGGTTTCGACTAC | YHALNPIGFDY | 11 | |
| b1F2-A-AAZ 5 | 1/12 | H31 | ACCTATTCCTTCTATTACTTTGCTTTCGACTAC | TYSFYYFAFDY | 11 | |
| b1F2-A-AAZ 7 | 1/12 | L50 | ACTGCCCACTTTTTTTCCTCTGCTCTCGACTAC | TAHFFSSALDY | 11 | |
| b1F2-A-AAZ 8 | 1/12 | L50 | CGGTCCCATACTTTCTATGTTGCTCTCGACTAC | RSHTFYVALDY | 11 | |
| b1F2-A-AAZ 9 | 1/12 | H31 | CATCACTACGACTATTCCTACGGTTTGGACTAC | HHYDYSYGLDY | 11 | |
| b1F2-A-AAZ 10 | 1/12 | H54 | TATAACCATGTCCCCTATTACGACTATAACTACCATCTTGCTCTGGACTAC | YNHVPYYDYNYHLALDY | 17 | |
| b1F2-A-AAZ 13 | 1/12 | L50 | CACCCCCACCTTTACTACCATGCTCTGGACTAC | HPHLYYHALDY | 11 | |
| b1F2-A-AAZ 14 | 1/12 | H54 | TACCCCTACTTTGCTAACTACGCTTTCGACTAC | YPYFANYAFDY | 11 | |
| b1F2-A-AAZ 16 | 1/12 | L50 | CGCAACTATCATTATGTTTATGGTCTCGACTAC | RNYHYVYGLDY | 11 | |
| b1F2-A-AAZ 17 | 1/12 | H31 | TATAACGAGTATCACTCTTTCGGTCTCGACTAC | YNEYHSFGLDY | 11 | |
| b1F2-A-AAZ 18 | 1/12 | H31 | TACGTGATTCATTATAACATTTTCCACCTCTTGGCTCTCGACTAC | YVIHYNIFHLLALDY | 15 | |
| b1F3-A-BS 1,3,5,8,9,11,12,13,14,15,16,17; b2F3-A-BS 3,4,7,8,11,13,14,15,18 | 21/30 | L28 | TACAACCCTTATTATGCTATGGACTAC | YNPYYAMDY | 9 | |
| b1F3-A-BS 7 | 1/30 | L28 | TATAATTCGGACTATCGTAATGGTATGGACTAC | YNSDYRNGMDY | 11 | |
| b1F3-A-BS 18 | 1/30 | L28 | TATAACGGTTATTACGCTATCGACTAC | YNGYYAIDY | 9 | |
| b2F3-A-BS 2,9,10,16,17 | 5/30 | L28 | TATGCTTACAACTATCGTGCTCTGGACTAC | YAYNYRALDY | 10 | |
| b2F3-A-BS 5 | 1/30 | L28 | TATTACCATAACTACGGTCTGGACTAC | YYHNYGLDY | 9 | Light chain Asp1 --> Gly |
| b2F3-A-BS 12 | 1/30 | L28 | TATTCTGACTATTATAGGGACGGTTTGGACTAC | YSDYYRDGLDY | 11 | |
| b1F3-O-BS 3,8,9,15,16,17; b2F3-O-BS 9,16 | 8/29 | L50 | TACGACCAGTATTTTTATGCTTTGGACTAC | YDQYFYALDY | 10 | |
| b1F3-O-BS 5, b2F3-O-BS 2,7,8,14,17 | 6/29 | L28 | TACCCTACCTACGCTGCCGATGGTATGGACTAC | YPTYAADGMDY | 11 | |
| b1F3-O-BS 6,7,11,13 | 4/29 | L50 | TACCTCCGCCCTTATTACGCTTTCGACTAC | YLRPYYAFDY | 10 | Sidlink Ser13 -> Asn |
| b1F3-O-BS 10,18 | 2/29 | L50 | CACGACCACAACTATTACGCTCTGGACTAC | HDHNYYALDY | 10 | |
| b1F3-O-BS 12; b2F3-O-BS 10,11,12,13,15 | 6/29 | L50 | TATGCCTCGGTTTATTATGCTCTCGACTAC | YASVYYALDY | 10 | |
| b1F3-O-BS 14 | 1/29 | L50 | TATCACATCACGTATTACGCTCTCGACTAC | YHITYYALDY | 10 | |
| b2F3-O-BS 6,18 | 2/29 | L50 | TATATTTCCAATTATTACGGTCTCGACTAC | YISNYYGLDY | 10 | |
| b1F4-A-AS 2,3,7,8,9,10,11,12,14,15,17 | 11/11 | H53* | CACTCCTATTATTACCTCGCTATCGACTAC | HSYYYLAIDY | 10 | Heavy chain ΔAla50 |

**Supplementary Table S5.** Coefficient of variation data for warhead swap experiment performed at 25 nM bCA.

| CV of cMyc+ population | | | | | | |
|---|---|---|---|---|---|---|
| Modification | b1F2 A-AAZ3 | b1F2 A-AAZ7 | b1F3 A-BS1 | b2F3 A-BS2 | b2F3 O-BS14 | b1F4 A-AS2 |
| Only AzF | 64.6 | 61.1 | 60.4 | 56.6 | 52.3 | 55.1 |
| AzF+4 | 72.3 | 76.7 | 80 | 76.8 | 81.2 | 61.2 |
| AzF+1 | 66.3 | 60.9 | 79.3 | 81.1 | 84.3 | 64.3 |
| AzF+3 | 57.3 | 57.4 | 65.6 | 67.8 | 65.7 | 64.6 |
| OPG+2 | 54.1 | 55.3 | 88.6 | 83.7 | 88.7 | 71.1 |
| Only OPG | 90.2 | 59.6 | 54.8 | 54.8 | 57.2 | 68.8 |
| CV of cMyc- population | | | | | | |
| Modification | b1F2 A-AAZ3 | b1F2 A-AAZ7 | b1F3 A-BS1 | b2F3 A-BS2 | b2F3 O-BS14 | b1F4 A-AS2 |
| Only AzF | 56.3 | 56.9 | 50.9 | 50.7 | 50.7 | 51.7 |
| AzF+4 | 60.8 | 60.8 | 61.6 | 65.8 | 58.4 | 60 |
| AzF+1 | 61.3 | 61.8 | 63.1 | 62.6 | 58.7 | 58.1 |
| AzF+3 | 54.9 | 64.3 | 59.2 | 58.9 | 56.6 | 54.7 |
| OPG+2 | 53.2 | 57.1 | 59.1 | 56.8 | 56.5 | 54.1 |
| Only OPG | 52.9 | 53.9 | 53.6 | 52.9 | 54.7 | 56 |

**Supplementary Table S6.** Coefficient of variation data for warhead swap experiment performed at 250 nM bCA.

| CV of cMyc+ population | | | | | | |
|---|---|---|---|---|---|---|
| Modification | b1F2 A-AAZ3 | b1F2 A-AAZ7 | b1F3 A-BS1 | b2F3 A-BS2 | b2F3 O-BS14 | b1F4 A-AS2 |
| Only AzF | 74.5 | 74.4 | 71.3 | 64.7 | 90.5 | 61.8 |
| AzF+4 | 79 | 80.3 | 77.6 | 73.4 | 92.2 | 70.4 |
| AzF+1 | 60.8 | 57.8 | 81.4 | 76 | 83.5 | 63.7 |
| AzF+3 | 81.1 | 60.9 | 71.7 | 76.7 | 75.5 | 73.1 |
| OPG+2 | 62.6 | 64.5 | 86.5 | 84.2 | 92.9 | 67 |
| Only OPG | 95.8 | 110 | 70.6 | 67.4 | 60.3 | 67.2 |
| CV of cMyc- population | | | | | | |
| Modification | b1F2 A-AAZ3 | b1F2 A-AAZ7 | b1F3 A-BS1 | b2F3 A-BS2 | b2F3 O-BS14 | b1F4 A-AS2 |
| Only AzF | 56.5 | 56.5 | 58.7 | 53.8 | 59.9 | 56.9 |
| AzF+4 | 75 | 68.3 | 66.8 | 71.8 | 136 | 59.2 |
| AzF+1 | 67.4 | 74.5 | 73.9 | 72.8 | 69.4 | 61.3 |
| AzF+3 | 61 | 57.7 | 72 | 65.4 | 79.3 | 66.5 |
| OPG+2 | 63.7 | 67.7 | 72.3 | 74 | 71.3 | 60.9 |
| Only OPG | 55.8 | 56 | 58.3 | 68.6 | 54.6 | 56 |

**Supplementary Table S7.** Conditions used for bCA inhibition assays.

| V (μL) | [scFv-Fc] (nM) | [4-NPA] (mM) | [bCA] (nM) | Note |
|---|---|---|---|---|
| 100 | 0 nM hybrid | 2 | 50 | |
| | 200 nM hybrid | | 50 | |
| | 200 scFv-Fc clicked w/o Cu | | 50 | scFv-Fc mixed with small molecule and all click chemistry reagents except for Cu |
| | 200 nM stock scFv-Fc | | 50 | scFv-Fc not subjected to CuAAC |
| | 200c hybrid w/o enzyme | | 0 | Control for hydrolyzation of 4-NPA from the hybrids |
| | 0 | | 0 | Negative control |
| | 200 nM SM | | 50 | Small molecule alone |

**Supplementary Table S8.** Labeling antibodies used for flow cytometry and western blotting.

| Target | Primary labeling | Secondary labeling |
|---|---|---|
| c-Myc tag | Chicken anti c-Myc (1:250) | Goat anti-Chicken Alexa Fluor 647 (1:500) (ThermoFisher) |
| Biotinylated bCA (FACS) | N/A | Mouse anti-biotin PE (1:50) (BioLegend) |
| Biotinylated bCA (flow cytometry analysis) | N/A | Streptavidin Alexa Fluor 488 (1:500) (ThermoFisher) |
| Clickable biotin probe | N/A | Streptavidin Alexa Fluor 488 (1:500) (ThermoFisher) |

**Supplementary Table S9.** Primers used to amplify fragments from the populations for Illumina paired end sequencing. Illumina partial adapter sequences are underlined in the primers. NNNNNN is the position of barcodes used for different populations, which are trimmed before sequence analysis.

| | |
|---|---|
| Frag 1 fwd | ACACTCTTTCCCTACACGACGCTCTTCCGATCTNNNNNNGGCGGAGGGTCGGCTAGC |
| Frag 1 rev | GACTGGAGTTCAGACGTGTGCTCTTCCGATCTGGCACCACTGCTACTGCCAC |
| Frag 2 fwd | ACACTCTTTCCCTACACGACGCTCTTCCGATCTNNNNNNGTGGCAGTAGCAGTGGTGCC |
| Frag 2 rev | GACTGGAGTTCAGACGTGTGCTCTTCCGATCTTGAGGAGACGGTGACCAGGGT |

**Supplementary Table S10.** Barcode sequences for the different populations.

| Population | Barcode Sequence |
|---|---|
| naïve Library | CTAAAT |
| b1F1-A-AA | GAGACA |
| b1F2-A-BS | ATCCGG |
| b2F2-A-BS | TATTAA |
| b1F2-O-BS | CAATCT |
| b2F2-O-BS | TGAGAC |
| b1F3-A-AS | TTGACG |

**Supplementary Table S11.** List of plasmids for scFv and suppression machineries.

| Plasmid name | Plasmid purpose | Backbone | Auxotrophic marker | Antibiotic marker |
|---|---|---|---|---|
| pCTcon2 bCA b1F2-A-AA3/ b1F2-A-AA7/ b1F3-A-BS1/ b1F3-O-BS5/ b1F4-A-AS2/ b2F3-A-BS2 | Yeast surface display | pCTcon2 | Trp | Amp marker |
| pCHA-FcSup-TAA bCA b1F2-A-AA3/ b1F2-A-AA7/ b1F3-A-BS1/ b1F3-O-BS5/ b1F4-A-AS2/ b2F3-A-BS2 | Protein secretion | pCHA-FcSup-TAA | Trp | Amp marker |
| pRS315-KanRmod-AcFRS | TAG codon suppression machinery: TyrAcFRS | pRS315 | Leu | Kan marker |

**Supplementary Table S12.** Other primers used in this work.

| Name | Sequence | Use |
|---|---|---|
| Con2seqfwd | GTTCCAGACTACGCTCTGCAG | Sequencing primers for scFvs in pCTcon2 plasmids |
| Con2seqrev | GATTTTGTTACATCTACACTGTT | |
| pCHA_scFv_NheI_fwd | CCATACGACGTTCCAGACTACGCTGCTAGCGACATACAGATGACTCAAAGTCCC | For cloning scFvs from the pCTcon2 plasmids into pCHA-FcSup-TAA plasmids |
| pCHA_scFv_XmaI_rev | TTTGTCGGAACTTTTAGGTTCTACCCCGGGTGAGGAGACGGTGACCAGGGTTCCTTG | |
| scFvFc_seq_FP | TGCCATTGGCCTTAGCTCAACCGG | Sequencing primers for scFv-Fcs in pCHA-FcSup-TAA plasmids |
| scFvFc_seq_RP | CGGCTTTGGCGGGAACAAAAAGACG | |

**Supplementary Table S13.** Sequences of the pCHA-FcSup plasmids. All the scFv sequences are upper-case and start at the same position in the plasmids. Backbone sequence is lower case and only listed once in the first plasmid sequence, for simplicity.

pCHA-FcSup-TAA bCA b1F2-A-AA3

gacgaaagggcctcgtgatacgcctatttttataggttaatgtcatgataataatggtttcttaggacggatcgcttgcctgtaacttacacgcgcctcgtatcttttaatgatggaataatttgg
gaatttactctgtgtttatttatttttatgttttgtatttggattttagaaagtaaataaagaaggtagaagagttacggaatgaagaaaaaaaaataaacaaaggtttaaaaaatttcaacaa
aaagcgtactttacatatatatttattagacaagaaaagcagattaaatagatatacattcgattaacgataagtaaaatgtaaaatcacaggattttcgtgtgtggtcttctacacagaca
agatgaaacaattcggcattaatacctgagagcaggaagagcaagataaaaggtagtatttgttggcgatcccccctagagtcttttacatcttcggaaaacaaaaactattttttctttaatt
tctttttttactttctatttttaatttatatatttatattaaaaaatttaaattataattatttttatagcacgtgatgaaaaggacccaggtggcacttttcggggaaatgtgcgcggaacccctatttgt
ttatttttctaaatacattcaaatatgtatccgctcatgagacaataaccctgataaatgcttcaataatattgaaaaaggaagagtatgagtattcaacatttccgtgtcgcccttattcccttttt
tgcggcattttgccttcctgtttttgctcacccagaaacgctggtgaaagtaaaagatgctgaagatcagttgggtgcacgagtgggttacatcgaactggatctcaacagcggtaagatc
cttgagagttttcgccccgaagaacgttttccaatgatgagcacttttaaagttctgctatgtggcgcggtattatcccgtattgacgccgggcaagagcaactcggtcgccgcatacacta
ttctcagaatgacttggttgagtactcaccagtcacagaaaagcatcttacggatggcatgacagtaagagaattatgcagtgctgccataaccatgagtgataacactgcggccaact
tacttctgacaacgatcggaggaccgaaggagctaaccgcttttttcacaacatgggggatcatgtaactcgccttgatcgttgggaaccggagctgaatgaagccataccaaacga
cgagcgtgacaccacgatgcctgtagcaatggcaacaacgttgcgcaaactattaactggcgaactacttactctagcttcccggcaacaattaatagactggatggaggcggataa
agttgcaggaccacttctgcgctcggcccttccggctggctggtttattgctgataaatctggagccggtgagcgtgggtctcgcggtatcattgcagcactggggccagatggtaagcc
ctcccgtatcgtagttatctacacgacgggcagtcaggcaactatggatgaacgaaatagacagatcgctgagataggtgcctcactgattaagcattggtaactgtcagaccaagttt
actcatatatactttagattgatttaaaacttcatttttaatttaaaaggatctaggtgaagatcctttttgataatctcatgaccaaaatcccttaacgtgagttttcgttccactgagcgtcagac
cccgtagaaaagatcaaaggatcttcttgagatcctttttttctgcgcgtaatctgctgcttgcaaacaaaaaaaccaccgctaccagcggtggtttgtttgccggatcaagagctaccaa
ctctttttccgaaggtaactggcttcagcagagcgcagataccaaatactgtccttctagtgtagccgtagttaggccaccacttcaagaactctgtagcaccgcctacatacctcgctctg
ctaatcctgttaccagtggctgctgccagtggcgataagtcgtgtcttaccgggttggactcaagacgatagttaccggataaggcgcagcggtcgggctgaacggggggttcgtgca
cacagcccagcttggagcgaacgacctacaccgaactgagatacctacagcgtgagcattgagaaagcgccacgcttcccgaagggagaaaggcggacaggtatccggtaag
cggcagggtcggaacaggagagcgcacgagggagcttccagggggaaacgcctggtatctttatagtcctgtcgggtttcgccacctctgacttgagcgtcgatttttgtgatgctcgtc
aggggggcggagcctatggaaaaacgccagcaacgcggcctttttacggttcctggccttttgctggccttttgctcacatgttctttcctgcgttatcccctgattctgtggataaccgtatta
ccgcctttgagtgagctgataccgctcgccgcagccgaacgaccgagcgcagcgagtcagtgagcgaggaagcggaagagcgcccaatacgcaaaccgcctctccccgcgcgt
tggccgattcattaatgcagctggcacgacaggtttcccgactggaaagcgggcagtgagcgcaacgcaattaatgtgagttacctcactcattaggcaccccaggctttacactttatg
cttccggctcctatgttgtgtggaattgtgagcggataacaatttcacacaggaaacagctatgaccatgattacgccaagctcggaattaaccctcactaaagggaacaaaagctggt
accgggcccccccctcgatcgaggtcgacggtatcgatgagcttgatatcgaattccttgaattttcaaaaattcttactttttttttggatggacgcaaagaagtttaataatcatattacatgg
cattaccaccatatacatatccatatacatatccatatctaatcttacttatatgttgtggaaatgtaaagagccccattatcttagcctaaaaaaaccttctctttggaactttcagtaatacgct
taactgctcattgctatattgaagtacggattagaagccgccgagcgggtgacagccctccgaaggaagactctcctccgtgcgtcctcgtcttcaccggtcgcgttcctgaaacgcag
atgtgcctcgcgccgcactgctccgaacaataaagattctacaatactagcttttatggttatgaagaggaaaaattggcagtaacctggccccacaaaccttcaaatgaacgaatcaa
attaacaaccataggatgataatgcgattagttttttagccttatttctggggtaattaatcagcgaagcgatgatttttgatctattaacagatatataaatgcaaaaactgcataaccacttt
aactaatactttcaacattttcggtttgtattacttcttattcaaatgtaataaaagtatcaacaaaaaattgttaatatacctctatactttaacgtcaaggagaaaaaaccccggatcatgaa
ggttttgattgtcttgttggctatcttcgctgctttgccattggccttagctcaaccggttatttctactaccgtcggttccgctgcagaaggctctttggacaagagagaagcttacccatacga
cgttccagactacgctgctagcGACATACAGATGACTCAAAGTCCCAGTTCACTATCTGCGTCTGTTGGTGATAGAGTCACCATTACGT
GTAGAGCTTCTCAGTCGATTAGCTCGTACTTGAATTGGTATCAACAGAAACCAGGGAAAGCTCCAAAGTTGCTGATCTATTAG
GCATCTAGCTTACAAAGTGGTGTACCTTCCAGGTTTTCAGGCTCAGGATCTGGAACTGATTTCACACTTACCATATCATCCTT
ACAACCGGAAGATTTCGCCACATATTACTGCCAACAATCCTACTCTACTCCACCTACATTTGGTGGTGGCACTAAAGTGGAG
ATTAAGGGTACTACTGCCGCTAGTGGTAGTAGTGGTGGCAGTAGCAGTGGTGCCGAGGTGCAATTGCTAGAATCAGGAGGT
GGTTTGGTACAACCTGGTGGTAGCTTAAGGTTGTCTTGTGCTGCTAGTGGATTCACGTTTAGTAGCTATGCCATGTCATGGG
TTAGACAAGCTCCAGGTAAAGGCTTAGAATGGGTTTCTGCGATATCTGGATCTGGTGGGTCAACTTACTATGCAGATTCCGT
CAAAGGCAGATTTACCATTTCCAGAGACAATTCGAAGAATACACTGTACCTTCAGATGAACTCGTTACGTGCAGAAGATACTG
CTGTTTACTACTGTGCTAAGTACCATGCTCTCAACCCGATCGGTTTCGACTACTGGGGCCAAGGAACCCTGGTCACCGTCTC
CTCAcccggggtagaacctaaaagttccgacaaaacccatacttgtccaccatgccctgctccagaattgttaggtggtcccagcgtctttttgttcccgccaaagccgaaagacac
gttgatgatcagtagaaccccggaagtaacatgtgttgttgtggacgtgagtcacgaggaccccgaagtaaagtttaattggtacgtggatggtgttgaagttcacaacgccaagacta
agcctagagaagaacagtacaattcaacatatcgtgttgttagcgtgttgactgttcttcaccaagattggcttaatggtaaggaatacaaatgcaaagtatccaataaagcacttccag
ctccgatagagaagacaatcagtaaagccaaaggtcagcccagagaacctcaagtttataccttgcctccgtctagagacgaattgaccaaaaatcaagtctccctgacttgcctagt
gaagggcttttacccatccgatatagcggtagaatgggaatcgaatggacaaccggaaaacaactataagactactccccctgtattggattctgatggatctttctttttgtatagcaaatt
gaccgttgataagtctaggtggcagcagggaaacgtcttctcatgtagtgttatgcacgaagccttacacaatcactacacacaaaagtcattatccttgtctccaggtaagagatcttaa
ggtggaggaggctctggtggaggcggtagcggaggcggagggtcgggatccacgcgtgcagaacaaaagcttatctccgaagaagacttgcaggaactgacaactatatgcga
gcaaatcccctcaccaactttagaatcgacgccgtactctttgtcaacgactactattttggccaacgggaaggcaatgcaaggagtttttgaatattacaaatcagtaacgtttgtcagta
attgcggttctcacccctcaacaactagcaaaggcagccccataaacacacagtatgtttttaactcgagcgaaatctgataacaacagtgtagatgtaacaaaatcgactttgttccc
actgtacttttagctcgtacaaaatacaatatacttttcatttctccgtaaacaacatgttttcccatgtaatatccttttctatttttcgttccgttaccaactttacacatactttatatagctattcactt
ctatacactaaaaaactaagacaattttaattttgctgcctgccatatttcaatttgttataaattcctataatttatcctattagtagctaaaaaaagatgaatgtgaatcgaatcctaagaaaa
ttccccatggtctaagcggcgccaccgcgggggggagctccaattcgccctatagtgagtcgtattacaattcactggccgtcgttttacaacgtcgtgactgggaaaaccctggcgttac
ccaacttaatcgccttgcagcacatccccctttcgccagctggcgtaatagcgaagaggcccgcaccgatcgcccttcccaacagttgcgcagcctgaatggcgaatggcgcgacg
cgccctgtagcggcgcattaagcgcggcgggtgtggtggttacgcgcagcgtgaccgctacacttgccagcgccctagcgcccgctcctttcgctttcttcccttcctttctcgccacgttc
gccggctttccccgtcaagctctaaatcgggggctccctttagggttccgatttagtgctttacggcacctcgaccccaaaaaacttgattagggtgatggttcacgtagtgggccatcgcc
ctgatagacggtttttcgccctttgacgttggagtccacgttctttaatagtggactcttgttccaaactggaacaacactcaaccctatctcggtctattcttttgatttataagggattttgccgat
ttcggcctattggttaaaaaatgagctgatttaacaaaaatttaacgcgaattttaacaaaatattaacgtttacaatttcctgatgcggtattttctccttacgcatctgtgcggtatttcacacc
gcaggcaagtgcacaaacaatacttaaataaatactactcagtaataacctatttcttagcatttttgacgaaatttgctattttgttagagtcttttacaccatttgtctccacacctccgcttac
atcaacaccaataacgccatttaatctaagcgcatcaccaacattttctggcgtcagtccaccagctaacataaaatgtaagctttcggggctctcttgccttccaacccagtcagaaatc
gagttccaatccaaaagttcacctgtcccacctgcttctgaatcaaacaagggaataaacgaatgaggtttctgtgaagctgcactgagtagtatgttgcagtcttttggaaatacgagtct
tttaataactggcaaaccgaggaactcttggtattcttgccacgactcatctccatgcagttggacgatatcaatgccgtaatcattgaccagagccaaaacatcctccttaggttgattac
gaaacacgccaaccaagtatttcggagtgcctgaactatttttatatgcttttacaagacttgaaattttccttgcaataaccgggtcaattgttctctttctattgggcacacatataataccca
gcaagtcagcatcggaatctagagcacattctgcggcctctgtgctctgcaagccgcaaactttcaccaatggaccagaactacctgtgaaattaataacagacatactccaagctgc
ctttgtgtgcttaatcacgtatactcacgtgctcaatagtcaccaatgccctccctcttggccctctcctttcttttttcgaccgaattaattcttaatcggcaaaaaaagaaaagctccggatc
aagattgtacgtaaggtgacaagctatttttcaataaagaatatcttccactactgccatctggcgtcataactgcaaagtacacatatattacgatgctgtctattaaatgcttcctatattata

tatatagtaatgtcgtttatggtgcactctcagtacaatctgctctgatgccgcatagttaagccagccccgacacccgccaacacccgctgacgcgccctgacgggcttgtctgctcccg
gcatccgcttacagacaagctgtgaccgtctccgggagctgcatgtgtcagaggttttcaccgtcatcaccgaaacgcgcga

pCHA-FcSup-TAA bCA b1F2-A-AA7

…GACATACAGATGACTCAAAGTCCCAGTTCACTATCTGCGTCTGTTGGTGATAGAGTCACCATTACGTGTAGAGCTTCTCAG
TCGATTAGCTCGTACTTGAATTGGTATCAACAGAAACCAGGGAAAGCTCCAAAGTTGCTGATCTATTAGGCATCTAGCTTACA
AAGTGGTGTACCTTCCAGGTTTTCAGGCTCAGGATCTGGAACTGATTTCACACTTACCATATCATCCTTACAACCGGAAGATT
TCGCCACATATTACTGCCAACAATCCTACTCTACTCCACCTACATTTGGTGGTGGCACTAAAGTGGAGATTAAGGGTACTACT
GCCGCTAGTGGTAGTAGTGGTGGCAGTAGCAGTGGTGCCGAGGTGCAATTGCTAGAATCAGGAGGTGGTTTGGTACAACCT
GGTGGTAGCTTAAGGTTGTCTTGTGCTGCTAGTGGATTCACGTTTAGTAGCTATGCCATGTCATGGGTTAGACAAGCTCCAG
GTAAAGGCTTAGAATGGGTTTCTGCGATATCTGGATCTGGTGGGTCAACTTACTATGCAGATTCCGTCAAAGGCAGATTTAC
CATTTCCAGAGACAATTCGAAGAATACACTGTACCTTCAGATGAACTCGTTACGTGCAGAAGATACTGCTGTTTACTACTGTG
CTAAGACTGCCCACTTTTTTTCCTCTGCTCTCGACTACTGGGGCCAAGGAACCCTGGTCACCGTCTCCTCA…

pCHA-FcSup-TAA bCA b1F3-A-BS1

…GACATACAGATGACTCAAAGTCCCAGTTCACTATCTGCGTCTGTTGGTGATAGAGTCACCATTACGTGTAGAGCTTCTCAG
TAGATTAGCTCGTACTTGAATTGGTATCAACAGAAACCAGGGAAAGCTCCAAAGTTGCTGATCTATGCAGCATCTAGCTTACA
AAGTGGTGTACCTTCCAGGTTTTCAGGCTCAGGATCTGGAACTGATTTCACACTTACCATATCATCCTTACAACCGGAAGATT
TCGCCACATATTACTGCCAACAATCCTACTCTACTCCACCTACATTTGGTGGTGGCACTAAAGTGGAGATTAAGGGTACTACT
GCCGCTAGTGGTAGTAGTGGTGGCAGTAGCAGTGGTGCCGAGGTGCAATTGCTAGAATCAGGAGGTGGTTTGGTACAACCT
GGTGGTAGCTTAAGGTTGTCTTGTGCTGCTAGTGGATTCACGTTTAGTAGCTATGCCATGTCATGGGTTAGACAAGCTCCAG
GTAAAGGCTTAGAATGGGTTTCTGCGATATCTGGATCTGGTGGGTCAACTTACTATGCAGATTCCGTCAAAGGCAGATTTAC
CATTTCCAGAGACAATTCGAAGAATACACTGTACCTTCAGATGAACTCGTTACGTGCAGAAGATACTGCTGTTTACTACTGTG
CTAAGTACAACCCTTATTATGCTATGGACTACTGGGGCCAAGGAACCCTGGTCACCGTCTCCTCA…

pCHA-FcSup-TAA bCA b1F3-O-BS5

…GACATACAGATGACTCAAAGTCCCAGTTCACTATCTGCGTCTGTTGGTGATAGAGTCACCATTACGTGTAGAGCTTCTCAG
TAGATTAGCTCGTACTTGAATTGGTATCAACAGAAACCAGGGAAAGCTCCAAAGTTGCTGATCTATGCAGCATCTAGCTTACA
AAGTGGTGTACCTTCCAGGTTTTCAGGCTCAGGATCTGGAACTGATTTCACACTTACCATATCATCCTTACAACCGGAAGATT
TCGCCACATATTACTGCCAACAATCCTACTCTACTCCACCTACATTTGGTGGTGGCACTAAAGTGGAGATTAAGGGTACTACT
GCCGCTAGTGGTAGTAGTGGTGGCAGTAGCAGTGGTGCCGAGGTGCAATTGCTAGAATCAGGAGGTGGTTTGGTACAACCT
GGTGGTAGCTTAAGGTTGTCTTGTGCTGCTAGTGGATTCACGTTTAGTAGCTATGCCATGTCATGGGTTAGACAAGCTCCAG
GTAAAGGCTTAGAATGGGTTTCTGCGATATCTGGATCTGGTGGGTCAACTTACTATGCAGATTCCGTCAAAGGCAGATTTAC
CATTTCCAGAGACAATTCGAAGAATACACTGTACCTTCAGATGAACTCGTTACGTGCAGAAGATACTGCTGTTTACTACTGTG
CTAAGTACCCTACCTACGCTGCCGATGGTATGGACTACTGGGGCCAAGGAACCCTGGTCACCGTCTCCTCA…

pCHA-FcSup-TAA bCA b1F4-A-AS2

…GACATACAGATGACTCAAAGTCCCAGTTCACTATCTGCGTCTGTTGGTGATAGAGTCACCATTACGTGTAGAGCTTCTCAG
TCGATTAGCTCGTACTTGAATTGGTATCAACAGAAACCAGGGAAAGCTCCAAAGTTGCTGATCTATGCAGCATCTAGCTTACA
AAGTGGTGTACCTTCCAGGTTTTCAGGCTCAGGATCTGGAACTGATTTCACACTTACCATATCATCCTTACAACCGGAAGATT
TCGCCACATATTACTGCCAACAATCCTACTCTACTCCACCTACATTTGGTGGTGGCACTAAAGTGGAGATTAAGGGTACTACT
GCCGCTAGTGGTAGTAGTGGTGGCAGTAGCAGTGGTGCCGAGGTGCAATTGCTAGAATCAGGAGGTGGTTTGGTACAACCT
GGTGGTAGCTTAAGGTTGTCTTGTGCTGCTAGTGGATTCACGTTTAGTAGCTATGCCATGTCATGGGTTAGACAAGCTCCAG
GTAAAGGCTTAGAATGGGTTTCTATATCTGGATAGGGTGGGTCAACTTACTATGCAGATTCCGTCAAAGGCAGATTTACCATT
TCCAGAGACAATTCGAAGAATACACTGTACCTTCAGATGAACTCGTTACGTGCAGAAGATACTGCTGTTTACTACTGTGCTAA
GCACTCCTATTATTACCTCGCTATCGACTACTGGGGCCAAGGAACCCTGGTCACCGTCTCCTCA…

pCHA-FcSup-TAA bCA b2F3-A-BS2

…GACATACAGATGACTCAAAGTCCCAGTTCACTATCTGCGTCTGTTGGTGATAGAGTCACCATTACGTGTAGAGCTTCTCAG
TAGATTAGCTCGTACTTGAATTGGTATCAACAGAAACCAGGGAAAGCTCCAAAGTTGCTGATCTATGCAGCATCTAGCTTACA
AAGTGGTGTACCTTCCAGGTTTTCAGGCTCAGGATCTGGAACTGATTTCACACTTACCATATCATCCTTACAACCGGAAGATT
TCGCCACATATTACTGCCAACAATCCTACTCTACTCCACCTACATTTGGTGGTGGCACTAAAGTGGAGATTAAGGGTACTACT
GCCGCTAGTGGTAGTAGTGGTGGCAGTAGCAGTGGTGCCGAGGTGCAATTGCTAGAATCAGGAGGTGGTTTGGTACAACCT
GGTGGTAGCTTAAGGTTGTCTTGTGCTGCTAGTGGATTCACGTTTAGTAGCTATGCCATGTCATGGGTTAGACAAGCTCCAG
GTAAAGGCTTAGAATGGGTTTCTGCGATATCTGGATCTGGTGGGTCAACTTACTATGCAGATTCCGTCAAAGGCAGATTTAC
CATTTCCAGAGACAATTCGAAGAATACACTGTACCTTCAGATGAACTCGTTACGTGCAGAAGATACTGCTGTTTACTACTGTG
CTAAGTATGCTTACAACTATCGTGCTCTGGACTACTGGGGCCAAGGAACCCTGGTCACCGTCTCCTCA…

**Supplementary Table S14.** Deep sequencing diversity coverage.

| | # of unique sequences | Total # of sequences | Ratio |
|---|---|---|---|
| Naïve library | 58083 | 64014 | 1 |
| b1F1-A-AA | 5617 | 288123 | 51 |
| b1F2-A-BS | 336 | 28470 | 85 |
| b2F2-A-BS | 213 | 24087 | 113 |
| b1F2-O-BS | 254 | 40376 | 159 |
| b2F2-O-BS | 240 | 23573 | 98 |
| b1F3-A-AS | 318 | 27099 | 85 |

### Materials and Methods

*Materials*

The *Saccharomyces cerevisiae* strain RJY100 was constructed as described previously.[1] The ncAA *p*-azido-L-phenylalanine was purchased from Chem-Impex International, Inc., and *p*-propargyloxyphenylalanine was purchased from Iris Biotech GmbH. All restriction enzymes used for cloning were from New England Biolabs. Primary and secondary antibodies used for flow cytometry labeling were purchased from Exalpha Biologicals (chicken anti-cMyc), BioLegend (anti-biotin PE), and Thermo Fisher Scientific (goat anti-chicken Alexa Fluor 647, goat anti-mouse Alexa Fluor 488, streptavidin Alexa Fluor 488). All PCR amplifications were performed with New England Biolabs Q5 DNA polymerase. Synthetic oligonucleotides for cloning and sequencing were purchased from Eurofins Genomics or IDT DNA Technologies. Sanger sequencing in this work was performed by Quintara Biosciences (Cambridge, MA). Illumina paired end deep sequencing was performed by GENEWIZ from Azenta. Mix and Go! kits from Zymo Research were used to prepare competent *E. coli*. Epoch Life Science GenCatch™ Plasmid DNA Mini-Prep Kits were used to isolate plasmid DNA from *E. coli*. Frozen-EZ Yeast Transformation II kits and Zymoprep DNA isolation kit from Zymo Research were used to prepare and transform competent yeast, and isolate plasmid DNA from yeast, respectively. Dynabeads™ Biotin Binder from Thermo Fisher Scientific were used for magnetic bead sorting. Penicillin-Streptomycin 100× from Corning was added to the yeast cultures for a final concentration of 100 IU/mL for Penicillin, and 100 µg/ml for Streptomycin. Bovine serum albumin (BSA) used for protein production was purchased from Proliant. Bovine carbonic anhydrase (bCA) was purchased from Sigma Aldrich (Cat. No: C2624-100MG). EZ-Link™ NHS-LC-Biotin, Zeba™ Spin Desalting Columns, 7K MWCO, 0.5 mL and Pierce™ Biotin Quantitation Kits were purchased from Thermo Scientific and used for antigen biotinylation. Biotin-PEG$_3$-Azide, Biotin-PEG$_4$-Alkyne, and THPTA were purchased from Click Chemistry Tools for use in CuAAC. (+)-Sodium L-ascorbate, aminoguanidine hydrochloride, copper sulfate pentahydrate, and DMSO were purchased from Sigma Aldrich for use in CuAAC. 4–12% Bis-Tris mini gels, SimplyBlue Safestain and iBlot™ Transfer Stacks were purchased from Thermo Scientific and used for SDS-PAGE and Western Blot analysis.

*Yeast surface hybrid construction using click chemistry*

The ncAA-containing scFv library was induced according to the protocol described previously in a yeast surface display protocol[2]. The library and the enriched populations were induced in the presence of 1 mM AzF (Chem-Impex International) or OPG (Iris Biotech GmbH).Changes to the scale of induction is as follows: for the first round of bead-based enrichments, 2 L of cells comprising the library at OD = 1 were induced in the presence of 1 mM AzF, and 1 L of cells comprising the library at OD = 1 were induced in the presence of OPG. For the second round of bead-based enrichment, 1 billion cells from b1-A-BS and b1-O-BS were induced in 100 mL each. For FACS enrichments, 5 mL of cells comprising the library at OD = 1 were induced.

Copper-catalyzed azide-alkyne cycloaddition (CuAAC) reactions on yeast surface were performed as described previously with some modifications specific to the cell number and small molecules used.[2-4] For the first round of sorting, 10 billion cells per condition were clicked in 25 mL reaction volume with each clickable sulfonamide (40 billion cells in total). Each small molecule was added to a final concentration of 1 mM, and the reaction was run for 4 h at room temperature. Cells were then washed 3 times with ice-cold PBSA to terminate the reaction and remove excess reagents. For the second round of sorting for b1-A-BS and b1-O-BS populations, 1 billion cells were clicked and washed under the same conditions as for round 1, scaling down to 2.5 mL total reaction volume. For FACS, 4 million cells from each track were subjected to click chemistry in a total volume of 250 µL for sorting. For small molecules **1**, **2**, and **3**, the reaction was performed with a final concentration of 1 mM small molecule (2.5 µL 100 mM stock solution) for 4 hours at room temperature. For small molecule **4** the reaction was performed with a final concentration of 0.1 mM small molecule (1.25 µL 20 mM stock solution) for 15 minutes at room temperature. This is based on our previous observation that longer reaction times or higher concentrations of **4** lead to more complete modification but loss of specific binding.[5] CuAAC reactions with biotin probes used a final concentration of 0.1 mM probe and proceeded for 15 minutes at room temperature..

*Hybrid library sorting*

Bead-based enrichments were performed according to previously described protocols[2]. Bovine carbonic anhydrase (bCA) (Sigma Aldrich, C2624-100MG, Lot# SLCD2277) was chemically biotinylated using previously described protocol to reach a 1-2 biotin/bCA ratio[6]. For FACS sorting we used a BioRad S3e cell sorter. Hybrid displaying cells were incubated with bCA for 1.5 h at room temperature and then washed with ice cold PBSA three times before 10 min secondary labeling on ice. Antibodies used for labeling and antibody dilutions are listed in **Supplementary Table S8.**

*Deep sequencing analysis of sorted population*

DNA from the populations was extracted via yeast miniprep. 1.25-2.5×$10^7$ cells from each population were used for yeast miniprep with Zymoprep Yeast Plasmid Miniprep kit and protocol (Zymo research). To sequence the scFvs with Illumina paired-end sequencing, PCR was performed to isolate two ~450 bp fragments from the scFv using primers with the partial adapter sequence specified for Genewiz Amplicon EZ sequencing (**Supplementary Table S9, S10**). The purified product was prepared according to the sample preparation instructions from Genewiz. Data was analyzed with Geneious Prime 2021.2.2 and visualized with GraphPad Prism. Sequence logos were generated with WebLogo[7]

Deep sequencing data sets were separately trimmed in Geneious Prime and exported to ScaffoldSeq[8] for family clustering and an analysis into site-specific amino acid enrichment across the CDR-H3 loop for each sorted population. For each sorted population, 100,000 sequences were analyzed using the following system settings and input parameters:

| ***ScaffoldSeq* System Settings** | |
|---|---|
| Sequence Similarity Threshold | 0.8 |
| Frequency Dampening Power | 0.5 |
| Maximum Sequence Count | 100,000 |
| Assay Background Filter | On |
| Pairwise Analysis | On |
| Filter Coefficient | 10 |

| ***ScaffoldSeq* Input Sequence Parameters** | |
|---|---|
| Gene Start Sequence | AAG |
| 5’ Anchor Sequence | TACTGT |
| 3’ Anchor Sequence | GGCCAA |
| DNA After Region | GAC |

Data related to site-wise amino acid frequencies, pairwise epistatic interactions, and sequence clusters (families) were output as .csv files. In addition to exporting these data tables for visualization in Python (see *Visualization of ScaffoldSeq outputs in Python*), information including number of unique clones and sequence frequencies in each sorting track were also tabulated. Because ScaffoldSeq can cluster CDR-H3 sequences based on their sequence similarity, individual families could be identified and grouped based on a defined sequence similarity threshold. We could then compare the highest-frequency sequences from each cluster and determine how distinct these families are to each other within a given sorting track. Heatmaps were then generated again in Python using the BLOSUM64 family distance scoring metric embedded within the ScaffoldSeq analysis (**Supplemental Figure S3**).

*Visualization of ScaffoldSeq outputs in Python*

Anaconda Navigator was used to run the web-based Python notebook jupyter. In Python, the *pandas* package was utilized to visualize tables output by *ScaffoldSeq.* The *seaborn* package was also used to generate heatmaps for both the site-wise amino acid enrichment and family distance scoring matrices.

*Sanger sequencing*
Individual clone DNA was isolated following previously described protocol[2]. All plasmids were sequence verified with sequencing performed at Quintara Biosciences. Sequencing primers are listed in **Supplementary Table S12.**

*Flow cytometric analysis for individual clone characterizations*
Analytical flow cytometry was performed as described in previous work.[3] To prepare cells for flow cytometry analysis, 2 million freshly induced cells of each sample were washed three times in PBSA (1× PBS, pH 7.4, with 0.1% w/v BSA) and then conjugated with alkyne-azide click chemistry. Cells are then labeled in 96 well V-bottom plates for flow cytometry. Biotinylated bovine carbonic anhydrase was incubated with the cells at room temperature for 1 hour along with chicken anti cMyc antibodies on an orbital shaker at 150 RPM. Following binding, all subsequent steps were performed on ice or in a centrifuge chilled to 4 °C. Cells were washed three times with ice-cold PBSA. Secondary labelling was performed on ice for 10 minutes in the dark with 3 washes in between. Samples were diluted in and washed twice with ice-cold PBSA before final resuspension in PBSA for flow cytometry. The labeling reagents and concentrations used for primary and secondary labelling are listed in **Supplementary Table S8**. Labeling reagents were prepared in PBSA. Flow cytometry was performed on an Attune NxT flow cytometer (Life Technologies) in the Tufts University Science and Technology Center, and data was processed using FlowJo™ software. 10,000 events were collected per sample on the flow cytometer. Samples were gated to isolate single cells and gated on c-Myc detection levels to isolate c-Myc-positive (sample) and c-Myc-negative (background) populations. The extent of reaction of the cell surface CuAAC is calculated based on the extent of biotin labeling on the cell surfaces of c-Myc-positive cells. The equations are listed below (Equation 1-3):

$$Extent\ of\ SM\ click\ reaction = 1 - Extent\ of\ Biotin\ click\ reaction \quad \text{(Equation 1)}$$

$$Extent\ of\ Biotin\ click\ reaction = \frac{MFI\ sample - MFI\ background}{MFI\ positive\ control - MFI\ background} \quad \text{(Equation 2)}$$

$$Extent\ of\ SM\ click\ reaction = 1 - \frac{MFI\ sample - MFI\ background}{MFI\ positive\ control - MFI\ background} \quad \text{(Equation 3)}$$

The errors are calculated based on the Coefficient of Variation(CV) values of the MFI values (Equation 4-7)

$$Numerator\ CV = \sqrt{CV\ of\ MFI\ sample^2 + CV\ of\ MFI\ background^2} \quad \text{(Equation 4)}$$

$$Demonimator\ CV = \sqrt{CV\ of\ MFI\ control^2 + CV\ of\ MFI\ background^2} \quad \text{(Equation 5)}$$

$$CV\ of\ final\ value = \sqrt{\frac{Numerator\ CV}{Numerator\ value}^2 + \frac{Denominator\ CV}{Denominator\ value}^2} \quad \text{(Equation 6)}$$

$$Propagated\ Error = Final\ value(extent\ of\ Biotin\ \ reaction) * CV\ of\ final\ value$$
(Equation 7)

Due to the high level of noise in samples acquired on the s3e sorter, the resulting CV values are larger than the median fluorescence intensity; this prevents calculation of propagated error.

*Yeast surface titration flow cytometry*

2 million freshly induced RJY100 cells displaying scFvs were conjugated with small molecules via CuAAC in 1.7 mL microcentrifuge tubes. After conjugations, cells were pelleted, washed three times with 1× PBSA, and then resuspended in 1 mL PBSA. To prepare for binding titration, 15,000 cells were added to wells of 96-well V-bottom plates. Biotinylated bCA were serially diluted, with a top concentration of 1 μM, seven subsequent 4-fold dilutions, and a final PBSA blank (0 μM bCA). The enzyme dilutions were added to the wells (ensuring >10× excess enzyme relative to the total number of displayed constructs in the well[9]) and incubated on the orbital shaker at 150 RPM at room temperature for 2 hours. Following labeling, flow cytometry data collection was performed as described in the *Flow cytometric analysis methods for individual clone characterizations* above, with the exception that 3,000 events were collected per sample instead of the usual 10,000 events. All titrations were performed in technical triplicates. We resuspended the cells and collected flow cytometry data one hybrid at a time instead of resuspending the whole plate (4 hybrids) at once to minimize signal loss over time after resuspension. Data analysis to obtain binding affinities ($K_D$) were done using steps like those outlined in section *Flow cytometric analysis methods for individual clone characterizations* above, with a few modifications. MFI levels for bCA detection within c-Myc-positive and c-Myc-negative populations was determined using FlowJo. Background correction was performed by subtracting MFI values of bCA detection within c-Myc-negative populations from MFI values of bCA detection within corresponding c-Myc-positive populations. The background corrected MFI data was normalized within GraphPad Prism (by dividing all MFI values by the MFI value for the highest bCA concentration). Then, using the “Receptor binding – Saturation binding” model and “One site -- Specific binding” equation, the data was fitted to enable determination of $K_D$ and 95% confidence intervals.

*Warhead swap methods*

To swap warheads installed in hybrids, cultures of cells displaying the six exemplary clones (b1F2-A-AAZ3, b1F2-A-AAZ7, b1F3-A-BS1, b2F3-A-BS2, b2F3-O-BS14, and b1F4-A-AS2) were each split in two cultures (12 total): one was induced with OPG and the other was induced with AzF. Four million cells were collected from each of the AzF-induced cultures and then split into four samples to react with one of the three alkyne-containing pharmacophores (**1**, **3**, **4**) or biotin probe, and two million cells from each of the OPG-induced cultures were split into 2 samples to react with the azide-containing pharmacophore (**2**) or biotin probe. The reactions were conducted in 96 well plate following the previously established protocol[5] to create a total of 24 hybrid-displaying cell samples and 12 cell samples displaying biotin-conjugated controls. Each hybrid-displaying samples were split into two halves, one half was used to determine the extent of reaction with 2-step click reaction along with the biotin-conjugated controls, and the other half was split in two to be tested for binding with either 25 nM or 250 nM bCA. Samples were incubated with bCA for 1 h and then the cells were washed and analyzed on an Attune NxT flow cytometer (Life Technologies). Binding data was collected as described in *Flow cytometric analysis for individual clone characterizations* and then converted into heat maps using Microsoft Excel.

*Production and purification of soluble scFv-Fc*
The scFv sequences were cloned into a previously established yeast secretion plasmid, pCHA-FcSup-TAA plasmid[6, 10] using NheI and XmaI restriction sites and Gibson Assembly. Sequences of scFv was amplified with PCR using the pCHA_scFvFwd and pCHA_scFvRev primers (**Supplementary Table S12)**. Each PCR-amplified insert was gel-purified, extracted, and then recombined with the pCHA-FcSup-TAA vector digested at the NheI and XmaI restriction sites using Gibson Assembly. All plasmids were sequence verified with sequencing performed at Quintara Biosciences (**Supplementary Table S13**). Zymo competent RJY100 yeast cells were transformed with the pCHA-FcSup-TAA plasmids (TRP marker) containing the scFv sequences as well as pRS-315 plasmids (LEU marker) containing the amber codon suppression machinery (aaRS/tRNA pair, TyrAcFRS for AzF and OPG).

ScFv-Fcs were secreted and purified according to previously established protocols[2, 6] with the following modifications. Yeast culture was expanded in SDSCAA −Trp −Leu −Ura media at 30 °C and then induced in YPG with 1 mM final concentration of ncAA and 0.1% w/v bovine serum albumin (Proliant Biologicals) at 20 °C, shaken at 275-300 rpm. After four days of incubation, the culture was spun down for 20 minutes at 3214 rcf and the supernatant was buffered with 10× PBS, pH 7.4 and sterile filtered with 0.2 µm bottle top filters. The scFv-Fc was purified with protein A resin (GenScript). The eluant was buffer exchanged using Amicon Ultra-15 centrifugal filter units (30 kDa molecular weight cut-off, Millipore Sigma) into PBS, pH 7.4 and concentrated. Protein concentrations were measured via the absorbance of 280 nm light on a Nano Drop One instrument (Thermo Fisher). Protein purity was determined via sodium dodecyl sulfate polyacrylamide gel electrophoresis (SDS-PAGE) without the use of PNGase F.

*ScFv-Fc click chemistry*
Click chemistry for soluble protein was performed using the same conditions as for reactions on the yeast surface, but using purified scFv-Fcs[2-4]. Reactions were performed in 1.7 mL microcentrifuge tubes, with 250 µL reaction volumes. To each tube was added 220 µL 1× PBS pH 7.4. For small molecules **1**, **2**, and **3**, the reaction was performed with 1 mM final concentration small molecule (2.5 µL of a 100 mM stock solution) for 2 hours at room temperature. For small molecule **4** the reaction was performed with 0.1 mM small molecule (1.25 µL of a 20 mM stock solution) for 15 minutes at room temperature as noted above. Biotin probes were added to a final concentration of 0.1 mM with reaction times of 15 minutes at room temperature. After small molecule conjugation reactions were complete, proteins were buffer exchanged to achieve a $10^7$-fold dilution of the reaction solution to terminate the reaction and remove excess small molecule. To evaluate whether scFv-Fcs affected bCA activity, ScFv-Fcs that had undergone reactions with all the click chemistry reagents except for $CuSO_4$ and

THPTA (Tris(3-hydroxypropyltriazolylmethyl)amine) were used as controls. Western blots were used to verify the presence of clickable ncAA using corresponding biotin probes, and following reactions to install small molecules, Western blots were used to estimate the extent of reaction. SDS-PAGE using 4–12% Bis-Tris gels was performed on all samples in duplicate gels to control for potential loss of protein during reactions. One gel was stained with Coomassie SimplyBlue SafeStain to confirm the presence of protein. Western blots were performed by transferring the second PAGE gel to a nitrocellulose membrane using an iBlot2 Dry Blotting System (Life Technologies). Transferred membranes were blocked with 5% w/v BSA in TBS + 0.1% v/v Tween20, following which they were probed with streptavidin Alexa Fluor 488 in a 1:1000 dilution in 5% w/v BSA in TBS + 0.1% v/v Tween20 for the presence of biotin.

*Carbonic anhydrase inhibition assay*

Carbonic anhydrase esterase activity was measured by monitoring the hydrolysis of 4-nitrophenyl acetate (4-NPA) to 4-nitrophenol and acetic acid in the presence of the enzyme. Bovine carbonic anhydrase (Sigma Aldrich, C2624-100MG, Lot# SLCD2277) at a final concentration of 50 nM was incubated with hybrids or control scFv-Fcs at various concentrations: 0 or 200 nM for the single concentration inhibition assay, and 0, 1, 10, 50, 100, 200 nM for the concentration response curve (details of conditions listed in **Supplementary Table S4**). After 1 hour shaking at room temperature 150 RPM in PBS, pH 7.4, 4-NPA was added to a final concentration of 2 mM right before the start of data acquisition. The 100 mM stock solution of 4-NPA in acetone was carefully diluted 5x in PBS, and then 10 µL of the dilution was added to the reaction in a final enzymatic assay volume of 100 µL containing 2 mM 4-NPA.The absorbance of 4-nitrophenol at 405 nm was measured with a SpectraMax i3x microplate reader every minute for 30 minutes at room temperature. To analyze the data, linear fits of the absorbance at 405 nM were performed in GraphPad Prism. Data in the initial 4 minutes are omitted due to low signal level and high noise level. After the initial 4 minutes, data points that are higher than the two readings before and after it in the curve are removed as outliers. The rest of the data points were linearly fit to obtain the reaction rates, and the rate of spontaneous hydrolyzation of 4-NPA without enzyme was subtracted from the enzymatic rates. The bCA rates are then normalized to the value of the condition where there is only bCA and substrate in the buffer. The bCA rate bar graphs and dose response curves are generated using GraphPad Prism.
